# GNC is a regulator of metabolic and productivity responses to elevated CO_2_ in *Arabidopsis thaliana*

**DOI:** 10.1101/2025.05.29.656866

**Authors:** Jennifer C. Quebedeaux, Kavya Kannan, Amy Marshall-Colón, Andrew D.B. Leakey

**Affiliations:** Department of Plant Biology, University of Illinois Urbana-Champaign, Urbana, IL 61801, USA; Carl R. Woese Institute for Genomic Biology, University of Illinois Urbana-Champaign, Urbana, IL 61801, USA; Department of Crop Sciences, University of Illinois Urbana-Champaign, Urbana, IL 61801, USA

**Keywords:** Arabidopsis, Elevated carbon dioxide, Photosynthesis, Gene expression

## Abstract

Despite established understanding of plant physiological responses to elevated [CO_2_], the underlying genes are poorly understood. Soybean transcriptomics previously identified a GATA transcription factor, involved in carbon and nitrogen metabolism, as responsive to elevated [CO_2_]. Supported by *in silico* modeling, we therefore hypothesized that this gene plays a previously unrecognized role in responding to elevated [CO_2_]. Wildtype and a T-DNA insertion line of *Arabidopsis thaliana* for GNC (GATA, Nitrate Inducible, Carbon Metabolism Involved) were grown under three treatments: sustained ambient [CO_2_], sustained elevated [CO_2_], and transfer from ambient to elevated [CO_2_], to assess changes in their physiology, biochemistry, and transcriptome. Photosynthetic and biomass responses to elevated [CO_2_] and transfer [CO_2_] in plants lacking GNC were significantly weaker than WT. A lag of 25-73 hrs in transcriptomic responses after transfer to elevated [CO_2_] was consistent with indirect sensing, presumably via sugar signals. The breakdown of the gene expression network around GNC was most pronounced in the transfer treatment and suggests targets for further study of interactions between elevated [CO_2_] and sulfur and nitrogen metabolism. This work provides a case study of a CO_2_-responsive transcription factor that may be a compelling target for adapting crops to future growing conditions after further characterization.

**Summary Statement:** A GATA transcription factor modulates plant metabolic and productivity responses to elevated CO_2_.

## Introduction

Elevated [CO_2_] stimulates leaf-level photosynthesis resulting in the production of more carbohydrates which can lead to greater biomass (Stitt, 1991; Leakey et al., 2009a). However, imbalance between the sizes of sources and sinks of carbon, sometimes driven by inadequate nitrogen availability, can prevent the potential gains in biomass production at elevated [CO_2_] from being met (Arp, 1991). When this happens, the plant senses accumulation of carbohydrates and acclimates by transcriptionally down-regulating photosynthesis (Krapp and Stitt, 1995; Koch, 1996; Jang et al., 1997; Moore et al., 1999). Elevated [CO_2_] can also be associated with changes in amino acids, structural components, and secondary metabolites (Penuelas and Estiarte, 1998; Ainsworth et al., 2006; Teng et al., 2006; Ekman et al., 2007; Li et al., 2008; Zinta et al., 2018). Additionally, transcriptional reprogramming to allow for greater respiratory capacity has been observed in soybean grown under field conditions (Leakey et al., 2009b).

A very limited number genes have been implicated in the control of these molecular and physiological responses to growth at elevated [CO_2_]. Hexokinase is a well-known glucose sensor modulating gene expression, but hexokinase-independent signaling and direct glucose and sucrose sensing can also occur (Jang et al., 1997; Rolland et al., 2002; Moore et al., 2003; Smith and Stitt, 2007; Hanson and Smeekens, 2009). Transcription factors more broadly involved in regulation of photosynthesis have been studied (Saibo et al., 2009; Zhang et al., 2016; Wang et al., 2017; Kannan et al., 2019). However, few genes have been found to mediate plant carbon-nitrogen balance, even outside the context of responses to elevated [CO_2_] (Sato et al., 2009; Kang et al., 2010; Sang et al., 2012; Aoyama et al., 2014; Para et al., 2014).

Soybean transcriptomics identified several differentially expressed genes in the response to elevated [CO_2_], including a member of the B-GATA transcription factor family (+24%) (Leakey et al., 2009b). There are approximately 30 members of the GATA transcription factor family in Arabidopsis (Reyes et al., 2004). The B-GATA subfamily can be split into members with LLM domains or HAN domains (Behringer and Schwechheimer, 2015). B-GATAs with a LLM domain are responsible for controlling germination, greening, senescence, and flowering time, while B-GATAs with a HAN domain are embryo and floral development regulators. Arabidopsis has six functionally redundant LLM domain B-GATAs. GNC (GATA, Nitrate Inducible, Carbon Metabolism Involved) and GNL (GNC-like) are the most studied of this subfamily. Several studies have suggested that the LLM domain B-GATA genes have overlapping, but distinct functions (Hudson et al., 2011; Ranftl et al., 2016; Xu et al., 2017). ChIP-seq identified 1,475 genes to be associated with GNC binding (Xu et al., 2017). GNC has been shown to upregulate hexose transporters, but does not appear to directly regulate hexokinase-dependent sugar signaling (Bi et al., 2005; Hudson et al., 2011). GNC also modulates chlorophyll biosynthesis and glutamate synthase (GLU1) expression, with knock-out mutants showing a reduction in chlorophyll content and chloroplast number as well as decreased expression of GLU1 (Bi et al., 2005; Hudson et al., 2011; Chiang et al., 2012; Bastakis et al., 2018). Interestingly, over-expression of GNC improved photosynthetic efficiency in *Arabidopsis* roots and over-expression in *Populus trichocarpa* (*poplar*) leaves showed higher photosynthetic capability (Ohnishi et al., 2018; An et al., 2020). Over-expression of GNC in poplar also improved nitrate uptake, remobilization, and assimilation (Shen et al., 2022). Many GATA factors have been identified in rice and soybean and have been implicated in light and nitrogen dependent gene regulation further indicating their centrality (Reyes et al., 2004; Hudson et al., 2013; Zhang et al., 2015; Kannan et al., 2019).

Most of what we know about how plants respond to CO_2_ is from long-term experiments where [CO_2_] is sustained over the entire growth period (Ainsworth and Long, 2021). This reflects the importance of understanding plant responses to elevated [CO_2_] in the natural context where elevated [CO_2_] will impact crop yields and global biogeochemical cycles that influence the progression of climate change (IPCC, 2021). Maybe because of this focus, far fewer studies have studied plants as they were transferred from ambient [CO_2_] to elevated [CO_2_]. This is despite transfer studies being a very common way to explore the transcriptional drivers of responses to other abiotic factors, including drought, temperature, light, and nitrogen supply (Kaplan et al., 2007; Talame et al., 2007; Usadel et al., 2008; Koini et al., 2009; Nakashima et al., 2009; Miller et al., 2010; Franklin et al., 2011; Oelze et al., 2012). Those transfer experiments that did study elevated [CO_2_] were predominantly done when the topic was first gaining popularity 20+ years ago before modern tools for genome-wide transcript abundance were available and a small number of genes had to be the focus. It is thought that transcriptional changes in photosynthetic processes can occur within a few hours of transfer to elevated CO_2_ resulting in an effect over a time period of three to seven days (Koch, 1996). However, most studies following plants shortly after a transfer from ambient [CO_2_] to elevated [CO_2_] have focused on biochemical and protein responses rather than on transcripts. A study of *Phaseolus vulgaris* (common bean) transferred three-week-old plants from low to high CO_2_, finding a decline in Rubisco activation state within one hour and lower Rubisco content by day five (Sage et al., 1989). *Lycopersicon esculentum* (tomato) transferred from ambient [CO_2_] to elevated [CO_2_] showed rapid downregulation of Rubisco small subunit and higher sucrose content one day after transfer (Van Oosten and Besford, 1994). However, this evidence in tomato is limited by its measurements on expanding leaves. Similar experiments have been conducted in *Arabidopsis* showing an increase in carbohydrates within four hours and decrease in Rubisco expression, content, and activity around six days (Cheng et al., 1998; Paul and Foyer, 2001). One study in *Solanum tuberosum* (potato) measured instantaneous photosynthesis, which was stimulated one hour after plants were transferred from ambient [CO_2_] to elevated [CO_2_] (Katny et al., 2005). Other experiments that were identified utilizing transfer to elevated [CO_2_] either measured changes after 10-30 days in the new treatment, combined [CO_2_] with confounding stress treatments (i.e., heat, drought, light) or exposed plants to super high [CO_2_] (Kaplan et al., 2012; Ge et al., 2018; Zinta et al., 2018; Gasparini et al., 2019; Torralbo et al., 2019). A physiological, biochemical, and transcriptional assessment of transferring plants from ambient [CO_2_] to elevated [CO_2_] could help us better understand how plants are responding to future conditions.

Greater understanding of genes controlling metabolic responses to elevated [CO_2_] could aid efforts to manipulate crops for enhanced performance under elevated [CO_2_] (Ainsworth et al., 2008; Kannan et al., 2019). As of yet, GNC has only been studied at ambient [CO_2_]. We hypothesize that GNC plays an important and previously unrecognized role in regulating metabolic and growth responses to elevated [CO_2_]. And, we predict that this will partly be manifested by altered transcriptional responses to elevated [CO_2_], which can reveal new information about the potential for the diverse pathways of plant metabolism to be reprogrammed when photoassimilate supply increases. Thus, wildtype (WT) and a T-DNA insertion line of *Arabidopsis thaliana* for GNC (*gnc*) were grown under three treatment combinations: sustained ambient [CO_2_], sustained elevated [CO_2_], and transient elevated [CO_2_] to assess changes in their physiology, biochemistry, and transcriptome. The focus of this study was how *gnc* and WT vary in their responses to different CO_2_ treatments.

## Methods

### Plant Material and Growth Conditions

A T-DNA insertion line of *Arabidopsis thaliana* for GNC (At5g56860, SALK_001778C) was acquired from the Arabidopsis Biological Resource Center (https://abrc.osu.edu/) and genotyped by PCR (Supplemental Table 1). Two experiments were conducted with wildtype (ecotype Columbia-0; WT) and mutant plants (*gnc*). The first experiment compared the physiological response of WT and *gnc* plants to transfer and elevated [CO_2_] while the second experiment compared their transcriptional response. For both experiments, seeds were cold treated for 48 hours and grown in 514 cm^3^ pots on LC1 Sunshine Mix (Sun Gro Horticulture, Agawam, MA, USA) mixed with 20% v/v small grain vermiculite. Two growth chambers (PGR14; Conviron, Winnipeg, Canada) were used to provide the following conditions: 10/14 hour day/night cycle at 21/18 °C, 70% relative humidity (RH), and 300 μmol m^-2^ s^-1^ of photosynthetically active radiation (PAR). Independent dataloggers (HOBO; Onset, Cape Code, MA, USA) were used to track chamber conditions. CO_2_ concentration was maintained at a beginning-of-the-21^st^ century ambient concentration of 370 ppm in one chamber and an elevated concentration of 750 ppm in the other chamber using custom retrofitted CO_2_ scrubbing and delivery systems, as described in Markelz et al. (2014a). Plants were watered once per week with 40% Long Ashton solution (6.0 mM NH_4_NO_3_) for the first four weeks of growth and every four days for the remainder of the experiment (Hewitt and Smith, 1975). Plants were rotated within chambers every other day and between chambers once a week to minimize chamber light variation and chamber bias. Replicate plants experienced either: (1) ambient [CO_2_] over the entire growth period; (2) elevated [CO_2_] over the entire growth period; or (3) ambient [CO_2_] until 30 days after germination (DAG), when they were transferred to elevated [CO_2_] one hour before the middle of the day. Transcriptomes were assessed in all treatments 30, 31 and 33 DAG, which corresponded to 1 hour, 25 hours, and 73 hours after the change in [CO_2_] in the transfer [CO_2_] treatment. Physiological, biochemical and biomass traits were assessed in all treatments 34 and 35 DAG.

### Leaf-level Physiology

All gas exchange was conducted using the LI-6800 gas exchange system equipped with a 2 cm^2^ leaf cuvette (LI-COR Biosciences, Lincoln, NE, USA). Photosynthesis-CO_2_ response (A/Ci) curves were measured (900 μmol m^-2^ s^-1^ PAR, 21°C, 70% RH, Flow= 500 μmol s^-1^) on the youngest fully expanded leaf 34 DAG in all treatments (after four days at elevated [CO_2_] in the transfer treatment) (n=4). Reference CO_2_ was controlled for each setpoint. The starting [CO_2_] corresponded to growth [CO_2_]: 370 ppm for ambient grown plants and 750 ppm for transfer and elevated grown plants. The CO_2_ concentration was then decreased stepwise, brought back to their respective starting concentration, and then increased. Curves were fit according to standard biochemical models of C_3_ photosynthesis and assuming infinite mesophyll conductance (Farquhar et al., 1980). Temperature corrected V_c,max_ and J_max_ were calculated as described by Bernacchi et al. (2001). Nighttime respiration was measured in the middle four hours of the night on the youngest fully expanded leaf 34 DAG (CO_2R_= 400 ppm, 18°C, 70% RH, Flow= 400 μmol s^-1^; n=5-8). Light response curves were measured on 35 DAG starting from high light and decreasing stepwise to zero (21°C, 70% RH, Flow= 500 μmol s^-1^, CO_2R_= 370 or 750 ppm; n=3-4).

Following gas exchange measurements, leaves were excised and scanned for leaf area. After respiration measurements, leaves were oven dried at 70 °C for calculation of specific leaf area (SLA; n=4-8). The dried leaves were ground and run on an elemental analyzer for determination of leaf nitrogen content (Costech, Valencia, CA, USA; n=8). After light response curves were measured, leaf disks (0.801 cm^2^) were collected and frozen for biochemical assessment. Extraction and quantification of chlorophyll was performed as described in Lichtenthaler and Wellburn (1983) using 96% v/v ethanol (n=4). Carbohydrates and starch were extracted and quantified as described in Ainsworth et al. (2006) (n=3-4). Whole plant aboveground biomass was harvested on 35 DAG for determination of dry biomass (n=7-8).

### Statistical Analysis of Physiological Traits

With the exception of light-response curves, all physiological, biochemical and biomass data were analyzed using a two-way analysis of variance (ANOVA) (PROC GLM, SAS 9.4; SAS Institute, Inc. Cary, NC, USA) considering both genotype and CO_2_ treatment as fixed effects. Differences were assessed by a post-hoc Tukey’s multiple comparison test. Proc mixed repeated measures ANOVA was used for light response curves.

### RNA Extraction

Samples from fully mature leaves of wildtype and *gnc* plants from all three treatments were taken for transcriptomic analysis (n=4) at midday on 30, 31, and 33 DAG and immediately frozen in liquid nitrogen. These sampling dates correspond to 1 hour, 25 hours, and 73 hours after exposure to elevated [CO_2_] began in the transfer [CO_2_] treatment. RNA samples were prepared according to manufacturer protocol using the Qiagen RNeasy Plant Mini Kit and included DNase digestion. RNA concentration and quality were determined using a NanoDrop One (Thermo Fisher Scientific, Waltham, MA, USA) and a Bioanalyzer (Agilent Technologies, Santa Clara, CA, USA).

### RT-qPCR

Expression of GNC was assessed by RT-qPCR using the Luna Universal One-Step RT-qPCR kit (New England BioLabs, Ipswich, MA, USA). The RT-qPCR experiment was performed by using a CFX Connect real-time PCR system (BioRad, Hercules, CA, USA). For each sample, three technical replicates were loaded in 96-well plates. The PCR mix consisted of a final volume of 20 μl containing Luna Universal One-Step Reaction Mix, Luna WarmStart RT Enzyme Mix, 0.4 μM forward and reverse primers, and <1 μg of template RNA. Reference primer sequences were taken from Czechowski et al. (2005) and GNC primer sequences were taken from Klermund et al. (2016) and confirmed via primer efficiency analysis (Supplemental Table 1). The amplification program was composed of a reverse transcription stage of 10 min at 55°C, a denaturation stage of 1 min at 95°C, and 45 amplification cycles with a 10 sec step at 95°C and a 30 sec step at 60°C. This was followed by a dissociation stage where the temperature was decreased to 65°C for 5 sec and gradually increased back to 95°C. Threshold cycle (Ct) values were provided by the CFX software (BioRad, Hercules, CA, USA) and the expression levels of genes of interest were determined using the method described by Schmittgen and Livak (2008).

### RNA Sequencing

Samples were sent to the Roy J. Carver Biotechnology Center at the University of Illinois for library preparation (Illumina TruSeq Stranded mRNA) and sequencing (HiSeq 4000; Illumina, San Diego, CA, USA). 100 bp single-end sequencing was completed using 4 lanes with a target of 20 million reads/sample. Fastq files were generated and demultiplexed using FASTQC (version 0.11.8) (Andrews, 2010).

### Sequence Alignment and Processing

Adapters were trimmed and MultiQC (version 1.6) was performed (Ewels et al., 2016). Average per-base read quality scores were over 30 in all samples indicating those reads were high in quality. Salmon (version 0.13.1) was used to quasi-map reads to the transcriptome and quantify the abundance of each transcript (Patro et al., 2017). The transcriptome was first indexed, then quasi-mapping was performed to map reads to transcriptome with additional arguments --seqBias and --gcBias to correct sequence-specific and GC content biases, -- numBootstraps=30 to compute bootstrap transcript abundance estimates and -- validateMappings and --recoverOrphans to help improve the accuracy of mappings. Gene-level counts were then estimated based on transcript-level counts using the “bias corrected counts without an offset” method from the tximport package. This method provides more accurate gene-level counts estimates and keeps multi-mapped reads in the analysis compared to traditional alignment-based method (Soneson et al., 2015).

The *Arabidopsis thaliana* transcriptome and annotation (Araport11 from Ensembl) were used for quasi-mapping and count generation. This transcriptome is derived from genome TAIR10. Coding and non-coding transcripts were quantified. Since the quasi-mapping step only uses transcript sequences, the annotation gtf file (Arabidopsis_thaliana.TAIR10.44.gtf) was solely used to generate a transcript-gene mapping table for obtaining gene-level counts. Longer gene names were pulled from the org.At.tair.db package from Bioconductor (release 3.9) (Huber et al., 2015).

Percentage of reads mapped to the transcriptome ranged from 0.7 to 97.7%. One sample failed based on this metric, and another was flagged for follow-up investigation while all the others scored above 95%. The small fraction of unmapped reads were discarded, while the number of remaining reads (range: 14.4 - 24.4 million per sample) were kept for statistical analysis. To account for differences in total number of reads and RNA composition, TMM (trimmed mean of M values) normalization in the edgeR package was performed to calculate a normalization factor for each sample (Robinson et al., 2010; Robinson and Oshlack, 2010).

While the Ensembl TAIR10 Annotation Araport11 gene models have a total of 32,309 genes, some of these might not have detectable expression. Setting the detection threshold at 1 cpm (counts per million) in at least 4 of the samples in the study, resulted in 13,558 genes being filtered out, leaving 18,751 genes to be analyzed for differential expression. These genes accounted for 99.95% of the total reads. After filtering, TMM normalization was performed again and normalized log2-based count per million values (logCPM) were calculated using edgeR’s cpm() function with prior.count = 3 to help stabilize fold-changes of extremely low expression genes.

Multidimensional scaling in the limma package was used as a QC step to check for outliers (Ritchie et al., 2015). The normalized logCPM values of the top 5,000 variable genes were chosen to construct multidimensional scaling plots. MDS clustering showed that whatever was causing the unusual characteristics of the sample initially flagged for low % of mapped reads was controlled for with the TMM normalization because it did not show up as an outlier from the rest of the samples until dimension 8, which only explains 2.1% of the overall variance. One additional sample was identified as an outlier based on MDS clustering and removed from all further analyses.

### Differential Expression

Proc t-test (SAS 9.4; SAS Institute, Inc. Cary, NC, USA) was used to test RT-qPCR measures of GNC expression. Bioinformatic analysis of transcriptome data was performed by HPC Bio at the University of Illinois. The read quality check and count generation was done using the *Biocluster* high-performance computing resource through the Computer Network Resource Group at the Carl R Woese Institute for Genomic Biology. All analyses from summation of counts to the gene level were done in R (R Core Team, 2017). To analyze differential expression, a 3-way ANOVA model was fit using limma trend. One sample was removed as it was deemed to be causing undue influence and adding error to the model. A 20% false discovery rate correction was applied and the pairwise comparisons from sampling timepoint three (corresponding to 73 hours after [CO_2_] change in the transfer [CO_2_] treatment) were used in further analyses.

### Gene Ontology Enrichment Analysis

Gene lists were separated into common and unique responses between WT and *gnc* and filtered by a log2FC > 1 and log2FC < -1. Fischer’s exact test and Bonferroni correction using PANTHER web tool was conducted to determine significantly over-represented biological processes (Thomas et al., 2021).

### Metabolism Mapping and Correlation Network Construction

All differentially expressed genes from the transfer experiment were categorized based on MapMan ontology (version 3.60RC1). Pearson’s correlations (cutoff = 0.98) were calculated in Cytoscape 3.9.1 and used to generate correlation networks. Networks were filtered to include only edges unique to each genotype. A subnetwork of WT was created using first neighbors as well as first and second neighbors (1 hop) of GNC. The KO network was then queried for the edges present in the WT network.

## Results

### GNC modulates biomass response to elevated [CO_2_]

Total above-ground biomass production was not significantly different in WT and *gnc* plants at ambient [CO_2_] (Fig. 1). In WT, biomass was significantly stimulated to a similar degree by both transfer [CO_2_] (43%) and elevated [CO_2_] (45%) relative to ambient [CO_2_]. In contrast, in *gnc* plants, biomass was not enhanced sufficiently by transfer [CO_2_] or elevated [CO_2_] treatments to be statistically resolvable.

**Fig. 1.**
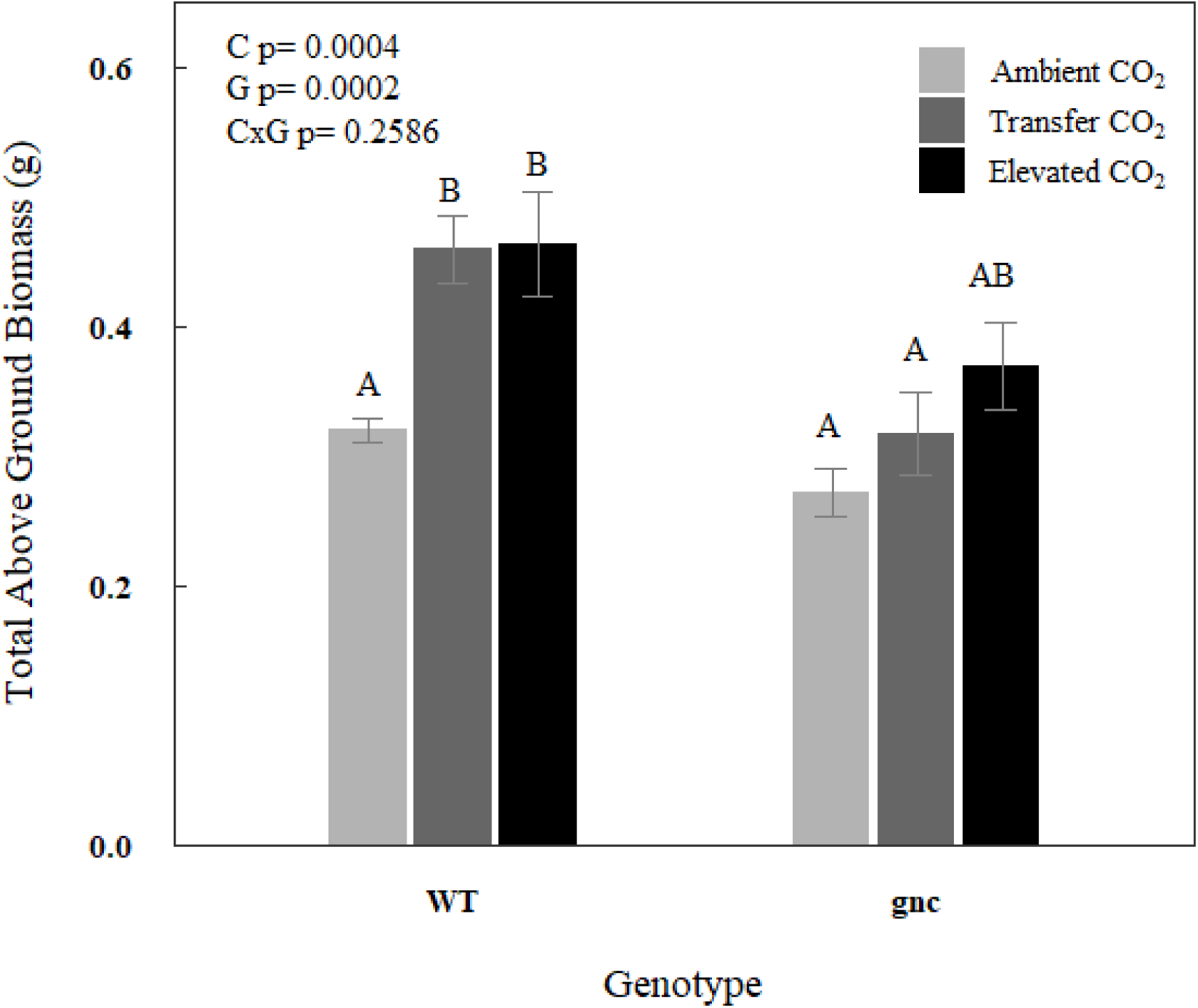
Biomass of WT and *gnc* plants under ambient [CO2], transfer [CO2], and elevated [CO2]. Bars show the mean value and error bars show the standard error. Letters indicate significance at *p* < 0.1. Abbreviations are as follows: CO2 Treatment (C) and Genotype (G).

*gnc plants had less of a stimulation of photosynthesis by transfer and elevated [CO_2_]*

There was a significant interactive effect of [CO_2_] treatment and genotype on photosynthetic CO_2_ uptake (p = 0.0017), which was consistent across the range of measurement light levels (CO_2_ x genotype x PAR p = 0.99; Fig. 2). There was no significant difference in the rate of photosynthetic CO_2_ uptake (*A*) in *gnc* compared to WT under ambient [CO_2_]. There was also no difference in *A* between plants that experienced long-term growth at elevated [CO_2_] versus short-term, transfer [CO_2_] treatment, regardless of whether they were WT or *gnc*. Therefore, the 2-way interaction effect was driven by the stimulation of *A* by both short- and long-term elevated [CO_2_] being less in *gnc* (+38% on average) than in WT (+64% on average). Consistent with the observed variation in *A* across treatments and genotypes, V_c,max_ and J_max_ were significantly lower in *gnc* than WT on average across the three [CO_2_] treatments (Fig. 3). And, there were no significant effects on V_c,max_ and J_max_ of elevated [CO_2_] or transfer [CO_2_] relative to ambient [CO_2_] in either *gnc* or WT. Similarly, nighttime respiration was significantly lower in *gnc* than WT on average across the three [CO_2_] treatments (Fig. 4). And, there were no significant effects on nighttime respiration of elevated [CO_2_] or transfer [CO_2_] relative to ambient [CO_2_].

**Fig. 2.**
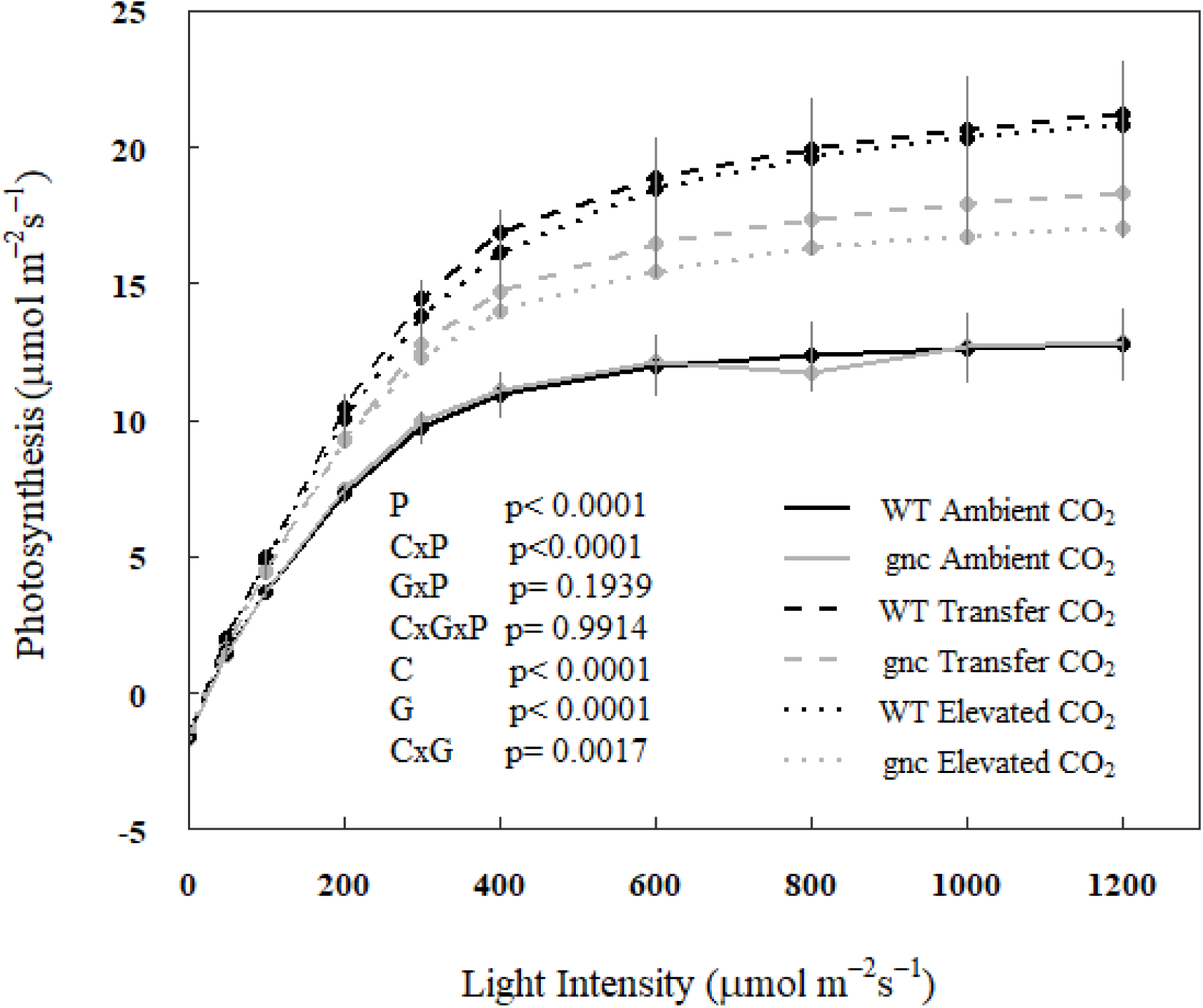
Light response curve of WT and *gnc* plants under ambient [CO2], transfer [CO2], and elevated [CO2]. Points show the mean value and error bars show the standard error. Abbreviations are as follows: Photosynthetically Active Radiation (P), CO2 Treatment (C), and Genotype (G).

**Fig. 3.**
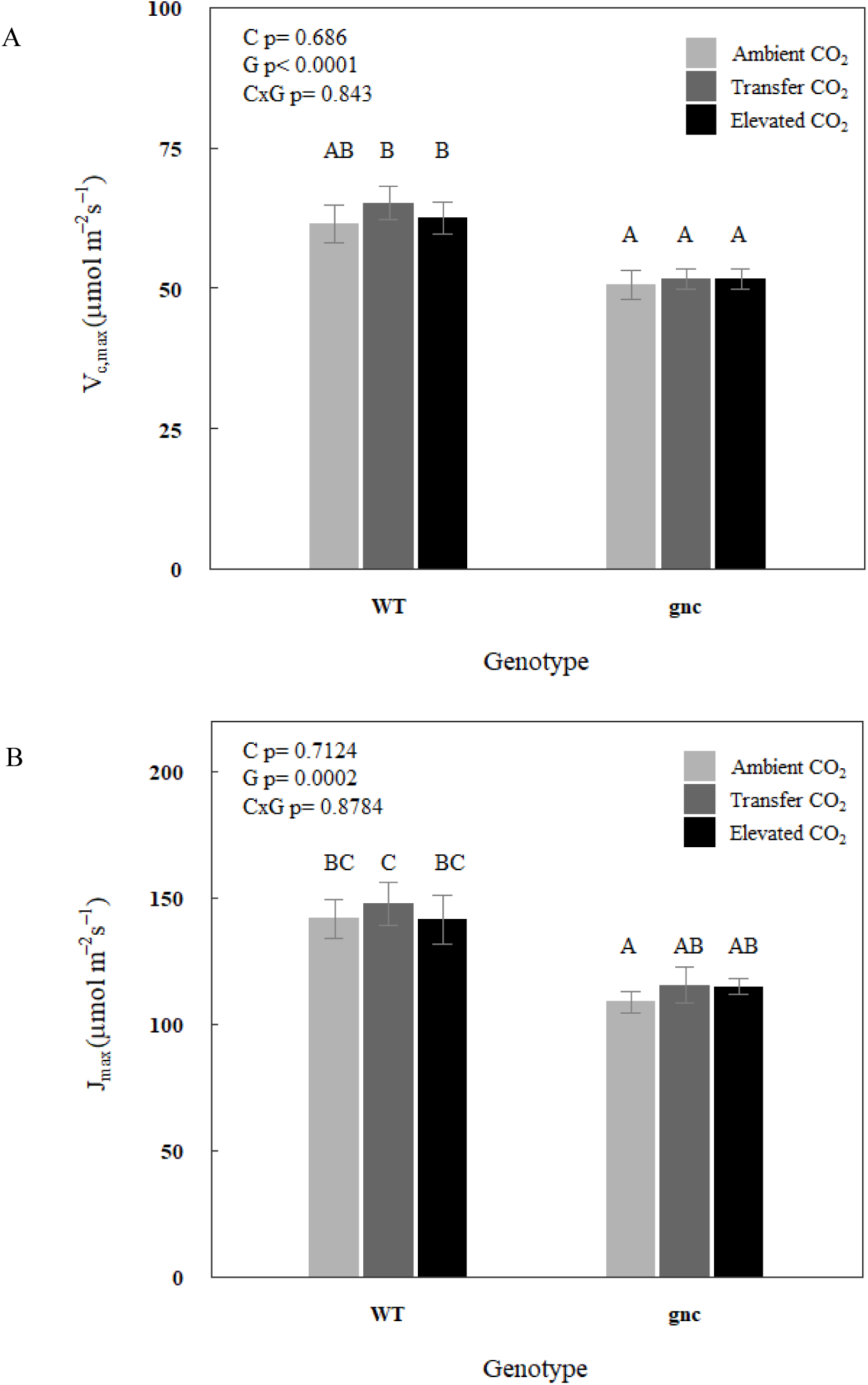
Gas exchange parameters maximum Rubisco carboxylation capacity (Vc,max) and maximum electron transport capacity (Jmax) of WT and *gnc* plants under ambient [CO2], transfer [CO2], and elevated [CO2]. Bars show the mean value and error bars show the standard error. Letters indicate significance at *p* < 0.1. Abbreviations are as follows: CO2 Treatment (C) and Genotype (G).

**Fig. 4.**
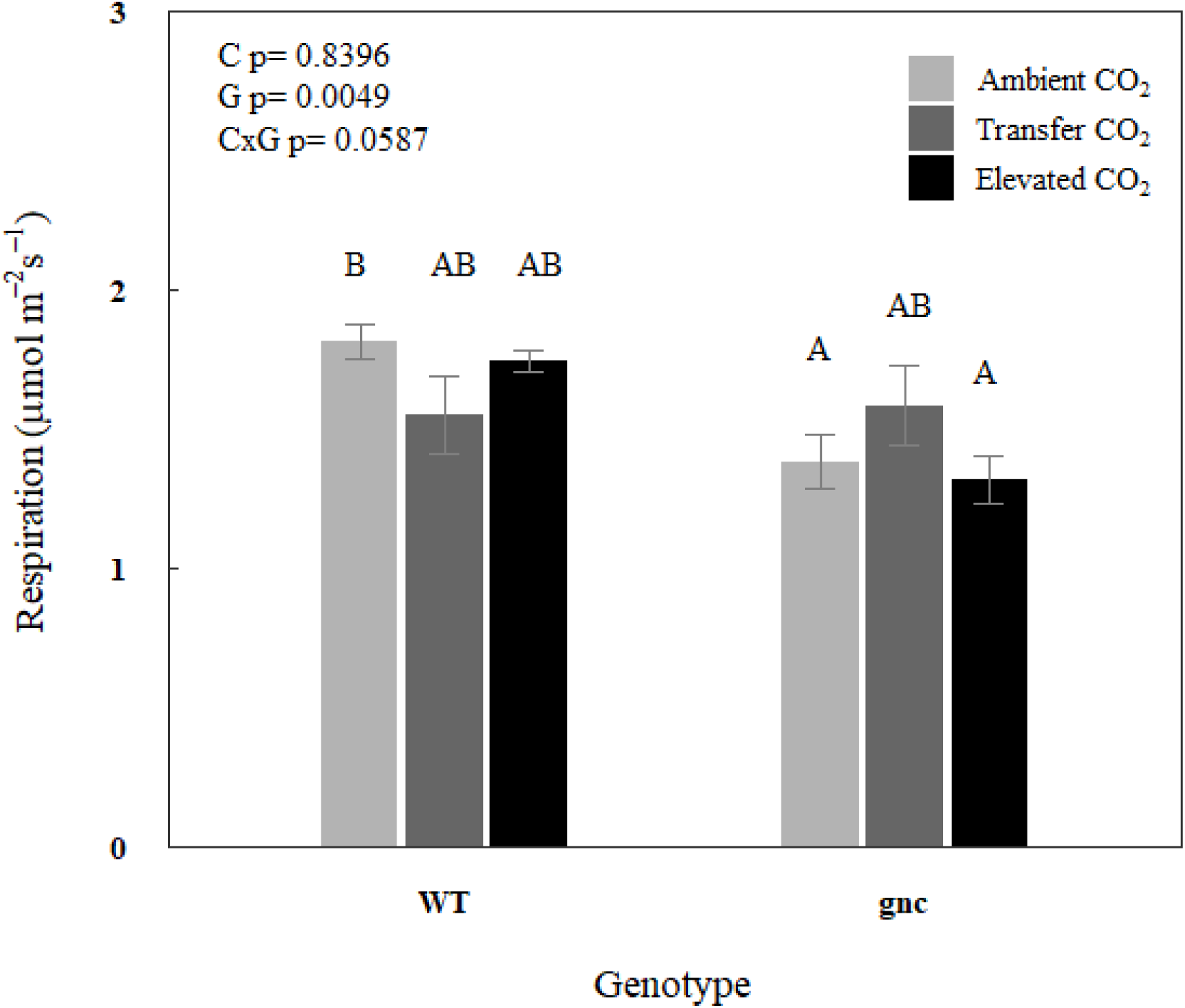
Respiration of WT and *gnc* plants under ambient [CO2], transfer [CO2], and elevated [CO2]. Bars show the mean value and error bars show the standard error. Letters indicate significance at *p* < 0.1. Abbreviations are as follows: CO2 Treatment (C) and Genotype (G).

### gnc showed greater changes in leaf N status

There were significant effects of genotype where, compared to WT, *gnc* had significantly: (1) lower leaf nitrogen (N) per unit area (Fig. 5A); (2) greater leaf N per unit mass (Fig. 5B); and (3) lower C:N ratio (Fig. 5C), on average across the three CO_2_ treatments. In addition, there were significant main effects of CO_2_ on all three measures of leaf N status (Fig. 5). Leaf N per unit mass responded equivalently in *gnc* and WT, being significantly lower in both transfer [CO_2_] and elevated [CO_2_] treatments relative to WT (Fig. 5B). Meanwhile, pairwise contrasts revealed genotype-specific responses of: (1) greater leaf N content per unit area in response to long-term elevated [CO_2_] in WT but not *gnc* (Fig. 5A); (2) greater C:N in response to the transfer [CO_2_] treatment in WT (+21%) but not *gnc* (Fig. 5C); and (3) greater C:N in response to the elevated [CO_2_] treatment in *gnc* (+21%) but not WT (Fig. 5C).

**Fig. 5.**
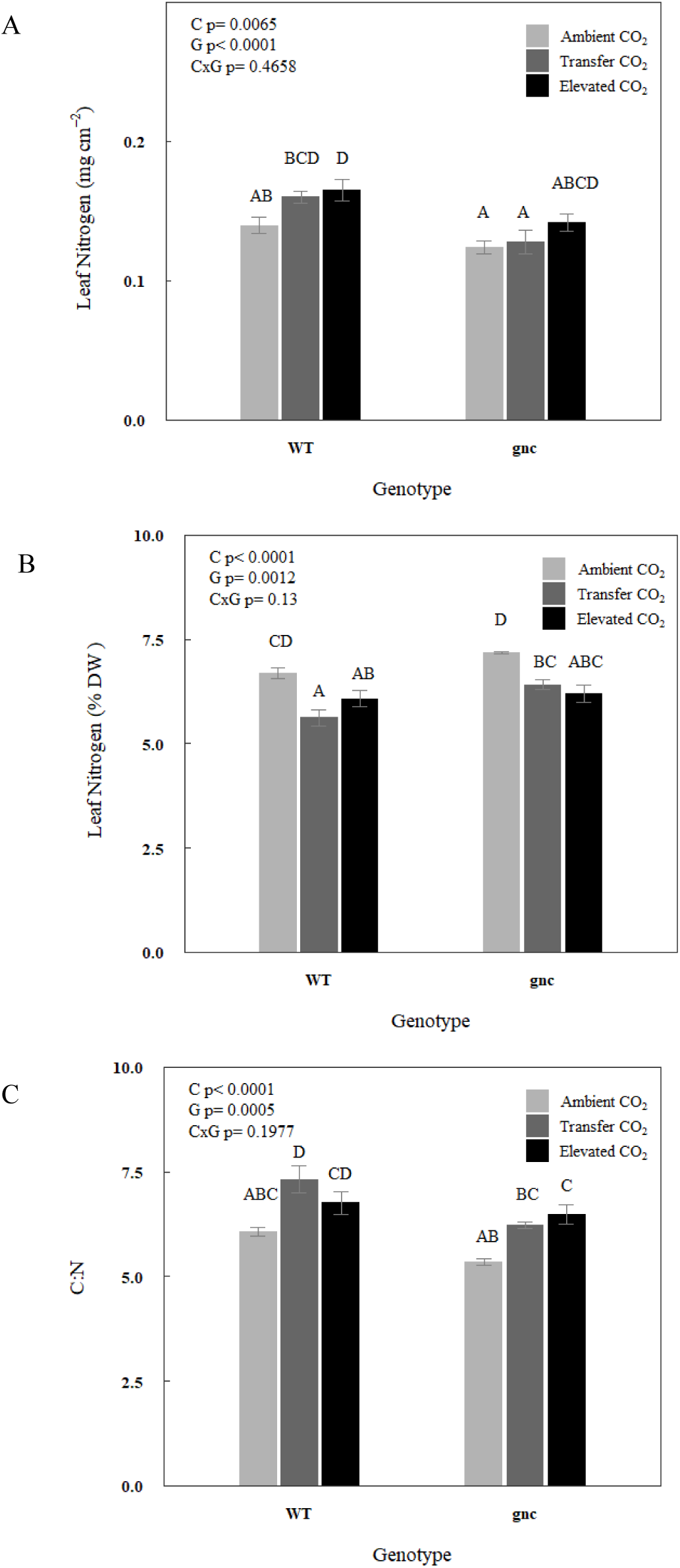
Leaf nitrogen composition A) per unit area, B) per unit mass, and C) as a ratio of carbon to nitrogen of WT and *gnc* plants under ambient [CO2], transfer [CO2], and elevated [CO2]. Bars show the mean value and error bars show the standard error. Letters indicate significance at *p* < 0.1. Abbreviations are as follows: CO2 Treatment (C) and Genotype (G).

### gnc showed altered patterns of response in SLA and lower chlorophyll content

Overall, leaf sugar and starch per unit area were both significantly greater when plants experienced some period of elevated [CO_2_] (Fig. 6 A-B). On average across CO_2_ treatments, *gnc* plants had significantly greater specific leaf area (SLA) than WT (Fig. 7). A significant decrease of SLA in WT was observed under transfer [CO_2_] (-22%) and elevated [CO_2_] (-23%) compared to ambient [CO_2_]. Meanwhile, a significant decrease of SLA in *gnc* was only observed under elevated [CO_2_] (-12%) compared to ambient [CO_2_]. Chlorophyll content trends lower in *gnc* plants at ambient [CO_2_] and is significantly lower in *gnc* compared to WT under transfer [CO_2_] and elevated [CO_2_] (Fig. 8).

**Fig. 6.**
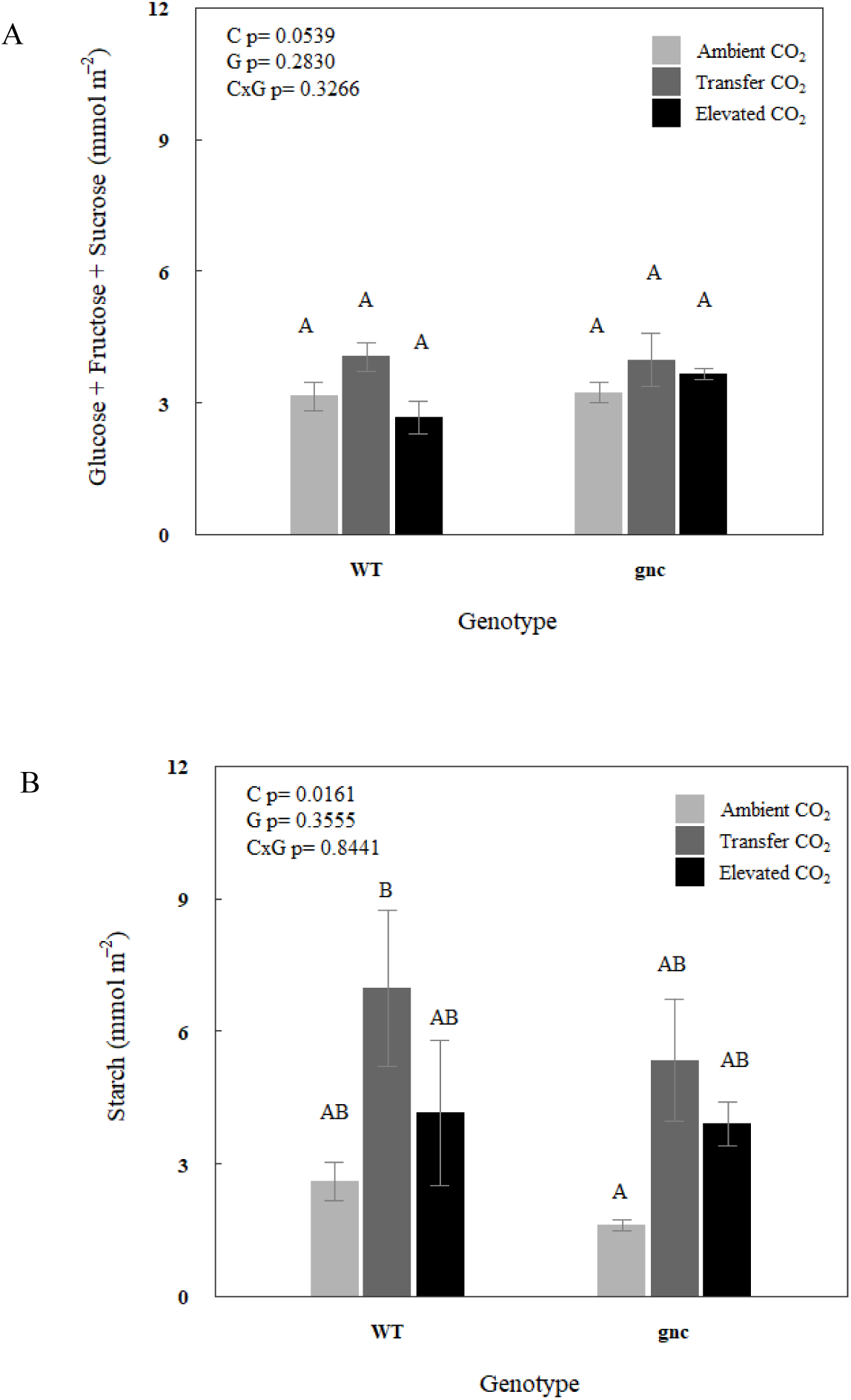
Total soluble sugar content (A) and starch content (B) of WT and *gnc* plants under ambient [CO2], transfer [CO2], and elevated [CO2]. Bars show the mean value and error bars show the standard error. Letters indicate significance at *p* < 0.1. Abbreviations are as follows: CO2 Treatment (C) and Genotype (G).

**Fig. 7.**
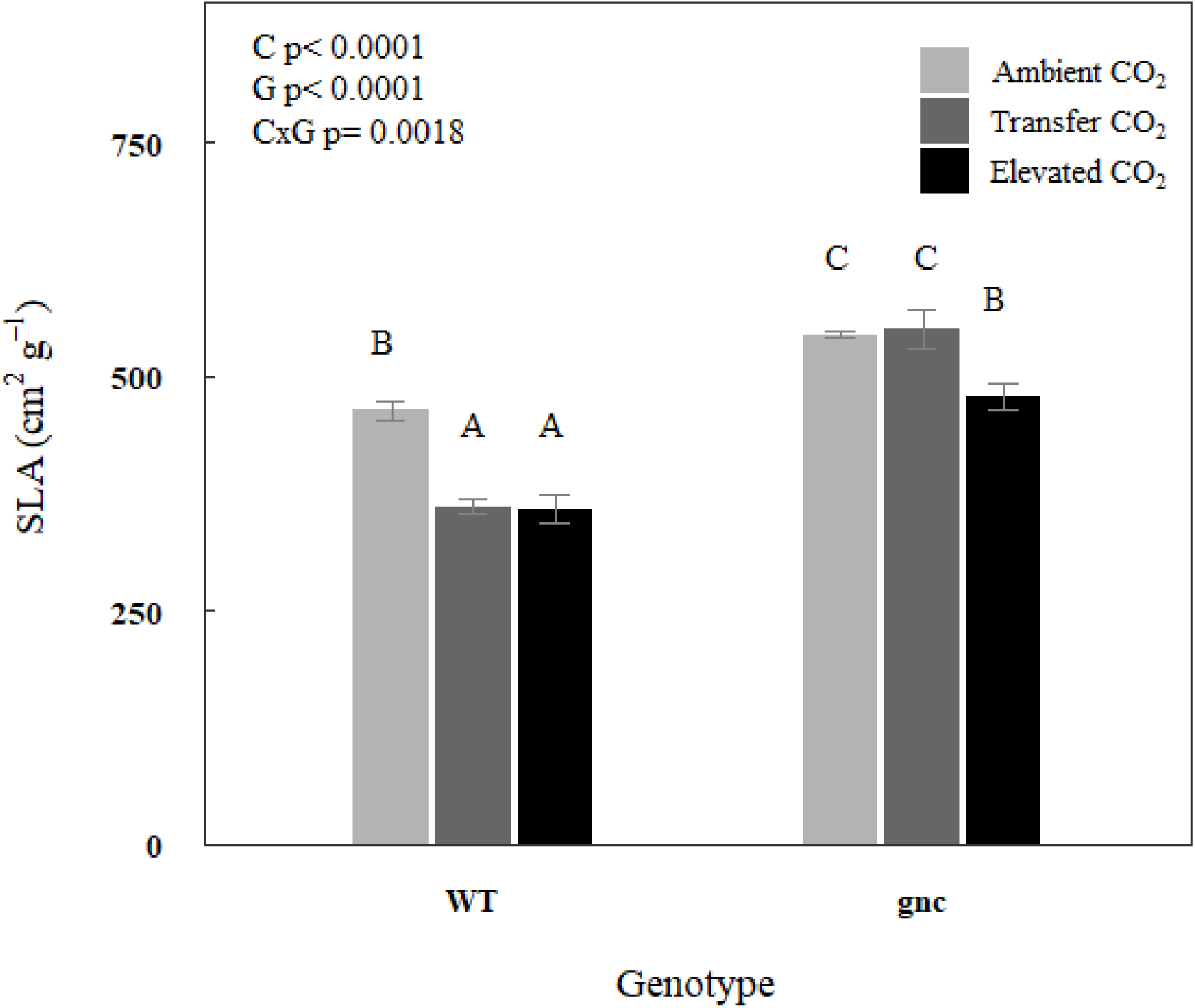
Specific leaf area (SLA) of WT and *gnc* plants under ambient [CO2], transfer [CO2], and elevated [CO2]. Bars show the mean value and error bars show the standard error. Letters indicate significance at *p* < 0.1. Letters indicate significance at *p* < 0.1. Abbreviations are as follows: CO2 Treatment (C) and Genotype (G).

**Fig. 8.**
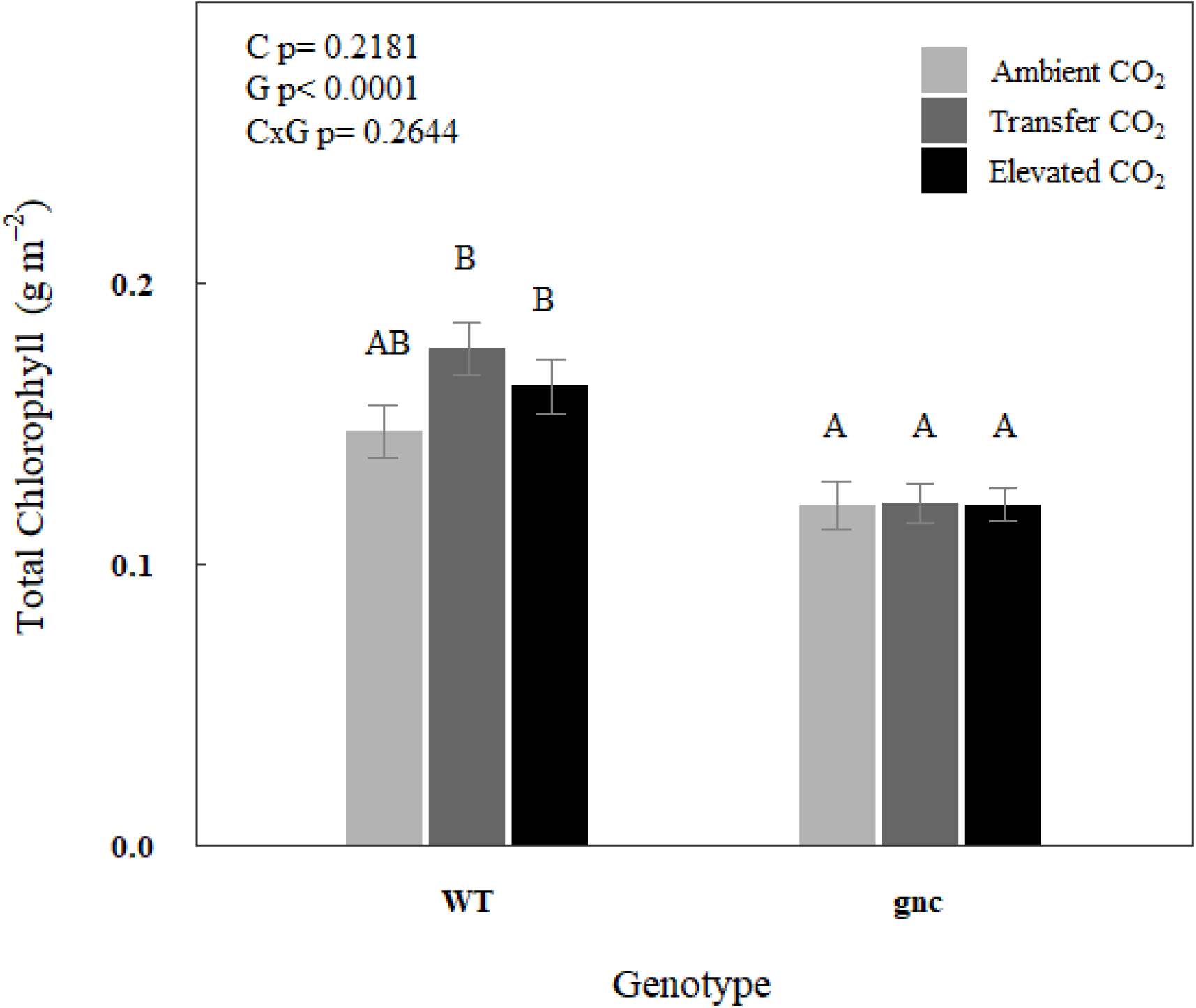
Chlorophyll content of WT and *gnc* plants under ambient [CO2], transfer [CO2], and elevated [CO2]. Bars show the mean value and error bars show the standard error. Letters indicate significance at *p* < 0.1. Letters indicate significance at *p* < 0.1. Abbreviations are as follows: CO2 Treatment (C) and Genotype (G).

### gnc shows similarity to previous transcriptomic reports

The expression of GNC was undetectable in *gnc* plants used for transcriptome analysis (Supplemental Fig. 1). Consistent with previous studies of *gnc* under standard growing conditions (Hudson et al., 2011), Glutamate synthase (GLU1; -0.401 log2FC) along with several genes involved in chlorophyll biosynthesis: HEMA1 (-0.3396 log2FC), PORB (-0.532 log2FC), PORC (-0.39 log2FC), and GUN4 (-0.464 log2FC) were all downregulated in *gnc* plants. PSII type I chlorophyll a/b binding protein, LHB1B1 (-0.95 log2FC), and light harvesting chlorophyll a/b binding protein, LHCB2.4 (-0.98 log2FC), were also downregulated. GNC-like (GNL) was the most upregulated gene in *gnc* plants (+3.359 log2FC).

### Lag and genotype effects in the transcriptional response to growth at elevated [CO_2_]

The abundance of many transcripts varied, on average, between CO_2_ treatments, genotypes and timepoints (Table 1). Of the 18,740 transcripts tested, only one transcript had a significant three-way interaction between [CO_2_], genotype, and time. Examining two-way interactions, the number of transcripts significant for the interaction between [CO_2_] and time was large (9,010 transcripts), while a moderate number of genes displayed an interaction between [CO_2_] and genotype (478 transcripts), and few genes were significant for the interaction between genotype and time (5 transcripts). Therefore, pairwise contrasts were performed for ambient [CO_2_] versus either elevated [CO_2_] or transfer [CO_2_] at each of the three timepoints and for each genotype. The long-term, sustained elevated [CO_2_] treatment altered the abundance of thousands of transcripts at all three timepoints in both genotypes. By contrast, the short-term, transfer [CO_2_] treatment triggered few to no changes in transcript abundance at the first or second timepoints. Nevertheless, by timepoint three (73 hours after transfer), the number of transcripts responding to the transfer [CO_2_] treatment (6,604 in WT, 5,107 in *gnc*) was similar to that responding to long-term growth at elevated [CO_2_] (4,840 in WT, 5,170 in *gnc*). Therefore, analyses focused on comparing *gnc* and WT in their responses to the CO_2_ treatments at the third timepoint.

**Table 1.**
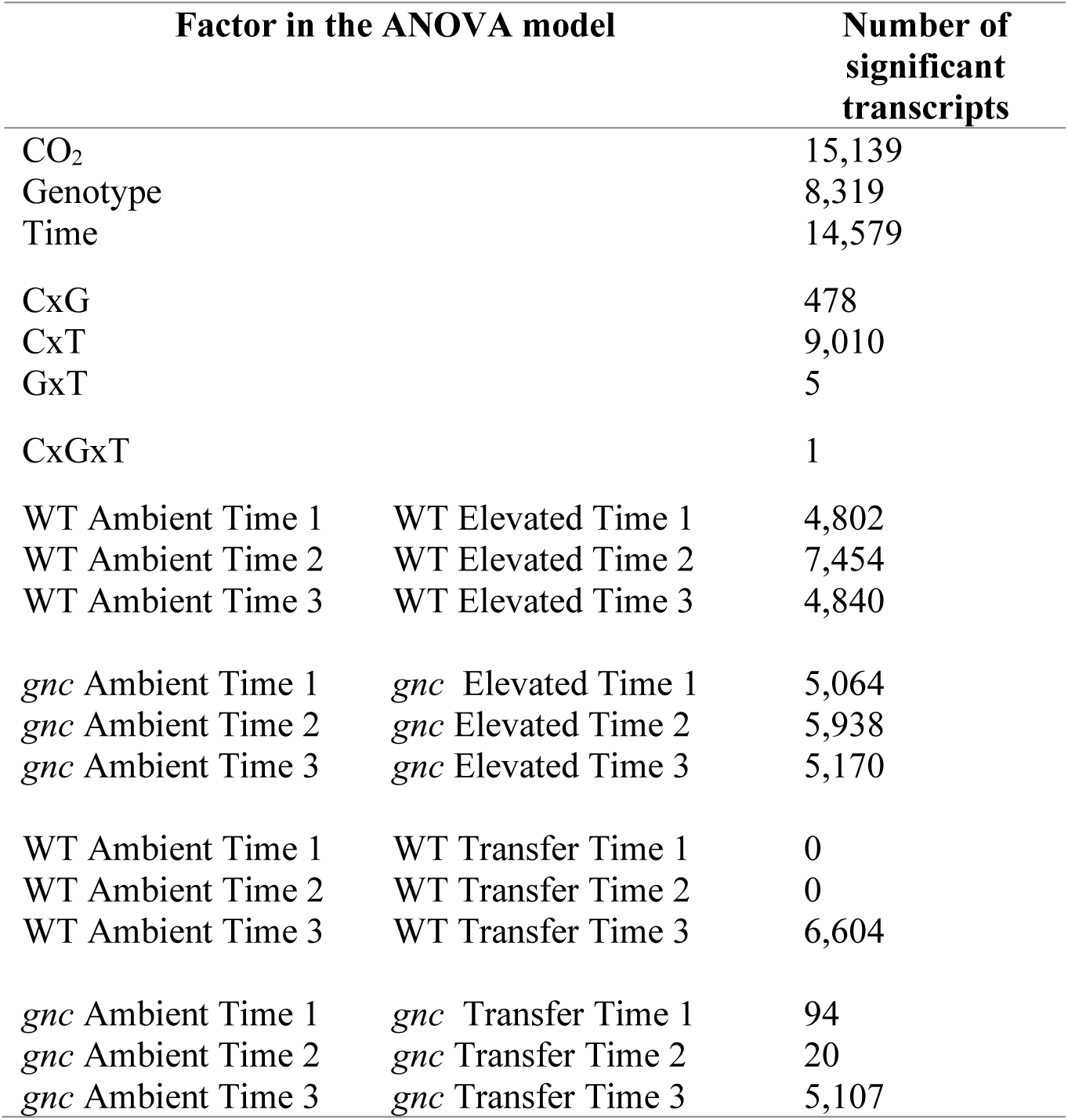
Number of transcripts responding significantly (FDR 20%) to each of the main effects and/or interactions in the ANOVA model of the 18,740 transcripts tested. Pairwise comparisons were conducted between ambient [CO2] and elevated [CO2] as well as ambient [CO2] and transfer [CO2] in WT and *gnc* at all three timepoints. Abbreviations are as follows: CO2 Treatment (C), Genotype (G), Time (T).

### gnc had a weaker response to transfer [CO_2_]

At timepoint 3, 45% of the differentially expressed (DE) genes in ambient [CO_2_] versus elevated [CO_2_] in *gnc* were also DE genes in WT (Fig. 9A). Meanwhile 60% of the DE genes in ambient [CO_2_] versus transfer [CO_2_] in *gnc* were also DE genes in WT (Fig. 9B). This snapshot of transcriptional responses in *gnc* and WT that were overlapping, but still contained a substantial number of distinct features, was reinforced by comparing the magnitude of changes in transcript abundance among the overlapping DE genes. When comparing ambient [CO_2_] versus elevated [CO_2_], DE genes common to *gnc* and WT had the strong tendency to respond with the same direction and magnitude of response (slope = 0.98, Fig. 10A). Meanwhile, when comparing ambient [CO_2_] versus transfer [CO_2_], DE genes common to both *gnc* and WT also tended to respond in the same direction, but more weakly in *gnc* than WT (slope = 0.85, Fig. 10B).

**Fig. 9.**
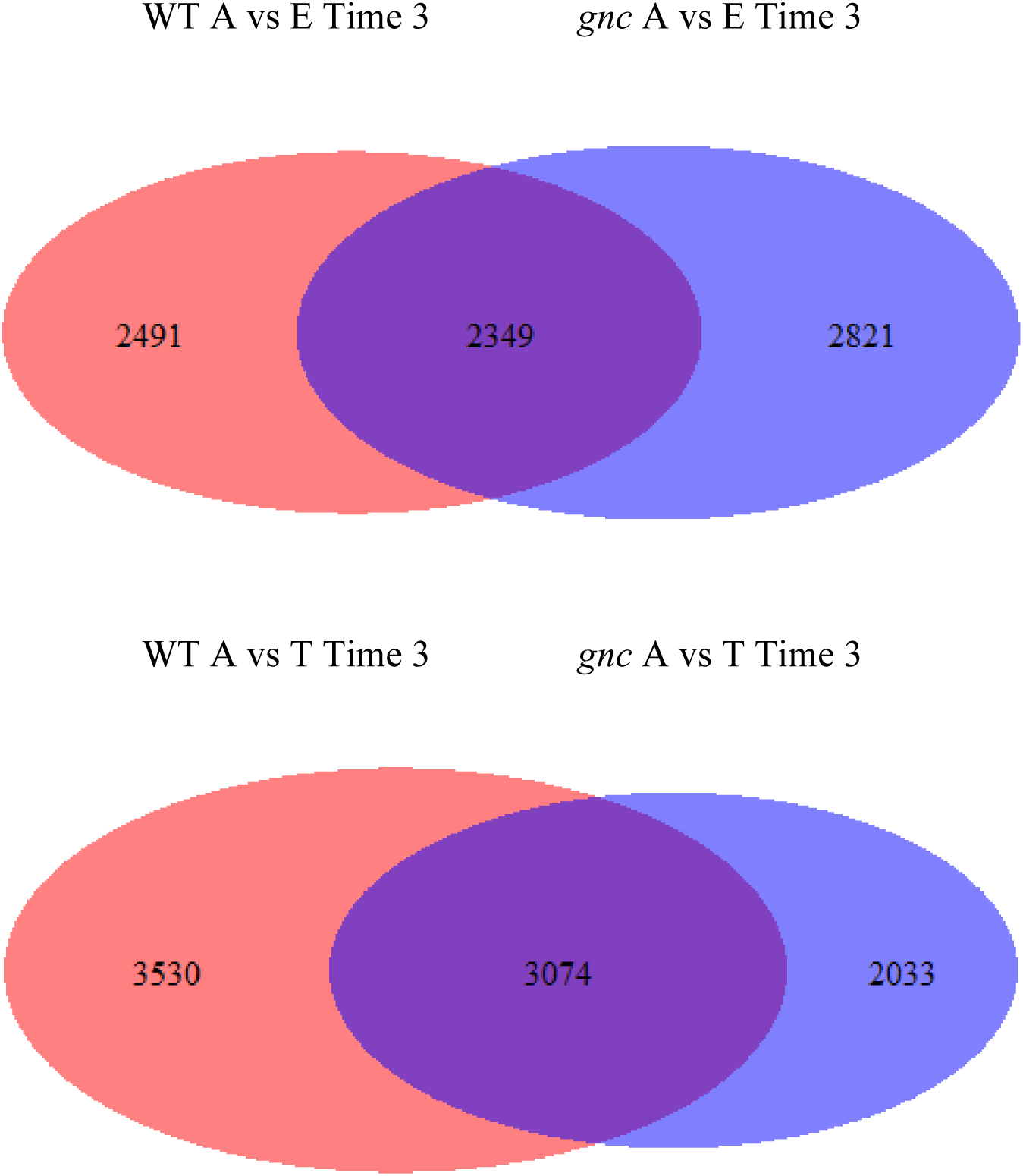
Venn diagrams depicting similarities and differences between WT and *gnc* in A) ambient [CO2] vs. elevated [CO2] as well as B) ambient [CO2] vs. transfer [CO2] at timepoint three. Diagram is scaled to indicate size of each component. Abbreviations are as follows: ambient [CO2] (A), elevated [CO2] (E), transfer [CO2] (T).

**Fig. 10.**
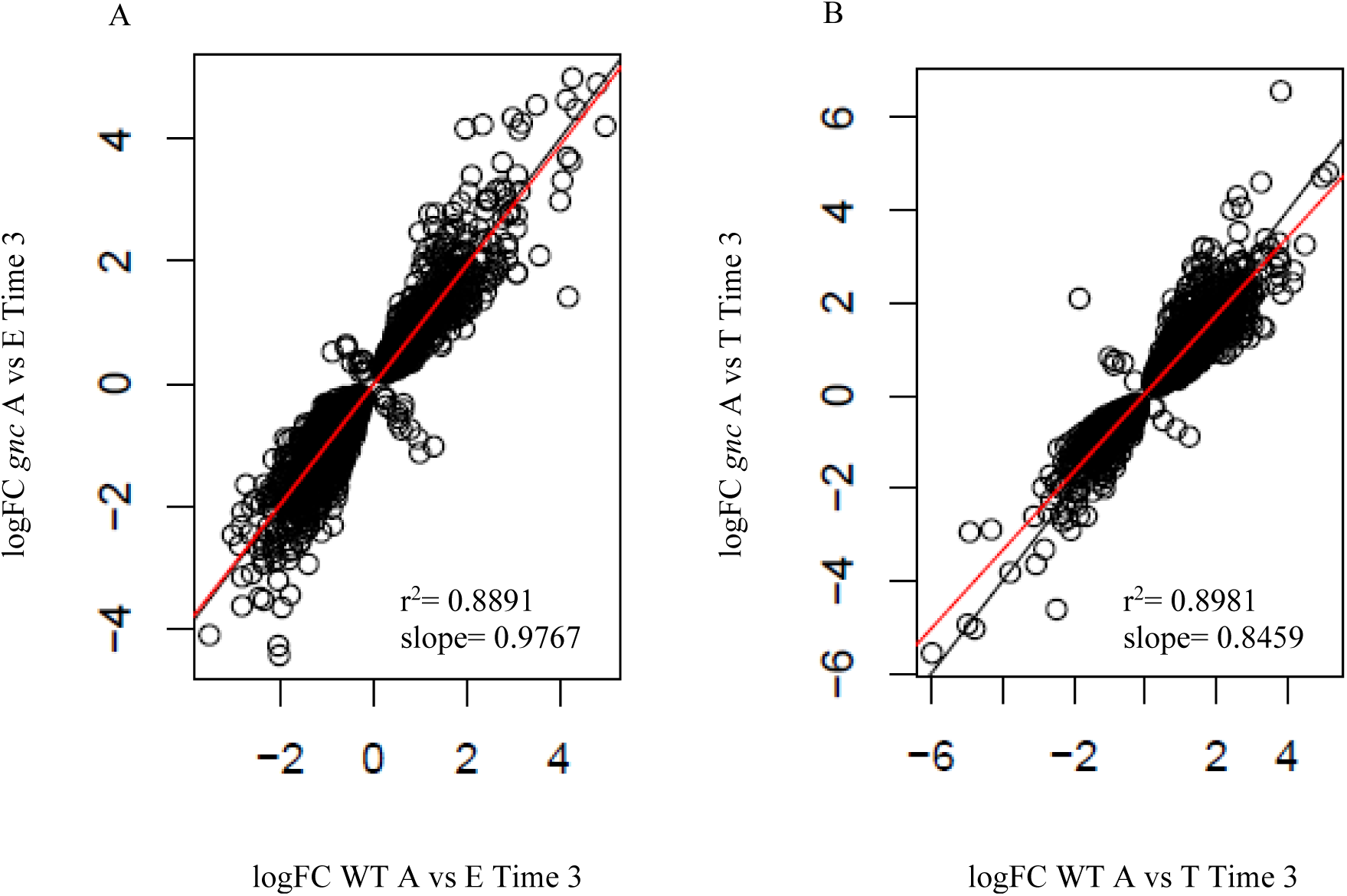
Regression plot of the relationship of genes responding significantly to elevated [CO2] (A) or transfer [CO2] (B) during timepoint 3. The red line is the line of best fit and the black line is a 1:1 line. In panel A, there were 834 genes responding in the positive direction for both genotype, 1,487 genes responding in the negative direction for both genotypes, and 28 genes responding in opposite directions. In panel B, there were 1,473 genes responding in the positive direction for both genotypes, 1,590 genes responding in the negative direction for both genotypes, and 11 genes responding in opposite directions. Abbreviations are as follows: ambient [CO2] (A), elevated [CO2] (E), transfer [CO2] (T).

### gnc plants display sulfur starvation transcriptional response when transferred from ambient [CO_2_] to elevated [CO_2_]

DE genes for ambient [CO_2_] versus elevated [CO_2_] at timepoint 3 were over-represented (p < 0.05) in multiple functional categories (Supplemental Table 2). In many cases, the responses were shared by *gnc* and WT, most notably: downregulation of sulfur compound biosynthesis, photosynthesis, and chlorophyll processes. Unique features of the response to elevated [CO_2_] in *gnc* included downregulation of cell wall modifications.

The functional categories that were over-represented with DE genes for ambient [CO_2_] versus transfer [CO_2_] at timepoint 3 that were common to *gnc* and WT included changes in several functional categories relating to lipid metabolism, fatty acid metabolism, and nitrogen metabolism (Supplemental Table 3). Unique features of the WT response to transfer [CO_2_] included downregulation of several functional categories related to RNA. Unique features of the *gnc* response included upregulation of sulfur starvation response and downregulation of glucosinolate biosynthesis.

### Distinct features of transcriptome responses for metabolic genes in gnc plants

Given that metabolism dominated the functional gene categories over-represented for DE genes, transcriptional responses to the CO_2_ treatments were mapped in greater detail onto metabolic pathways (Figs 11-14). Focusing first on responses to elevated [CO_2_] versus ambient [CO_2_] that were shared by *gnc* and WT, genes related to photosynthesis, carbonic anhydrases, and tetrapyrrole synthesis were downregulated (Fig. 11 and 12). Notably, this included a downregulation of the Rubisco small subunit (rbcS). Nitrogen metabolism genes also responded in both genotypes, with lower transcript abundance at elevated [CO_2_] of amino acid synthesis genes as well as key N assimilation genes, including nitrate reductase (NIA1), glutamine synthetase (GS2 and GLN1.2), and glutamate synthase (GLU1 and GLT1). There was a corresponding general increase in transcript abundance for genes involved in amino acid degradation in *gnc* and WT. There was modestly increased abundance of transcripts at elevated [CO_2_] in both genotypes for many respiratory genes, particularly in the tricarboxylic acid cycle and mitochondrial electron transport. Lipid degradation genes were generally upregulated at elevated [CO_2_] compared to ambient [CO_2_] in both genotypes; whereas cell wall degradation genes were generally downregulated. In secondary metabolism, elevated [CO_2_] led to down-regulation of a number of pathways in both genotypes, most notably terpene metabolism and S-miscellaneous.

**Fig. 11.**
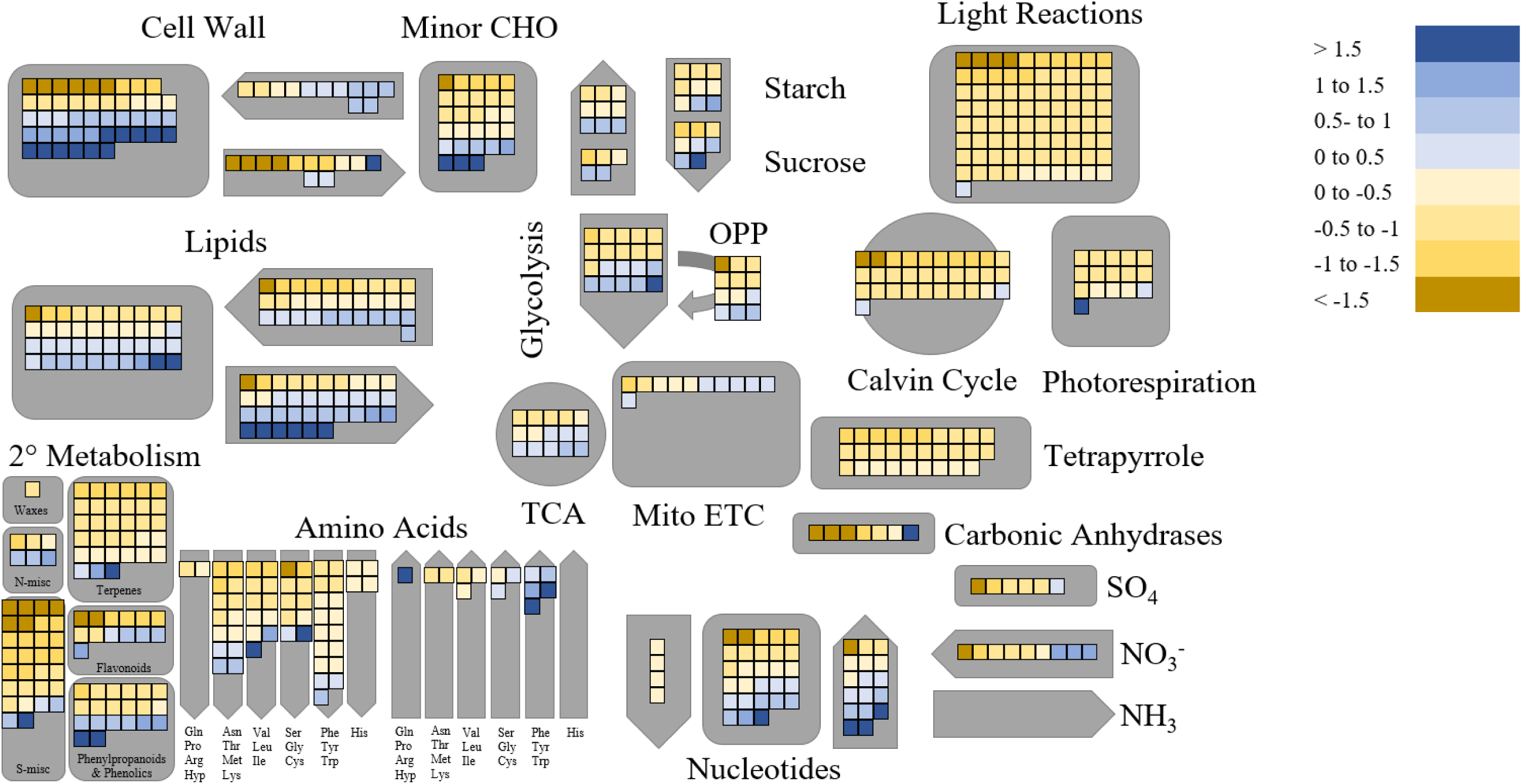
Transcript abundance in central metabolism pathways of WT response to elevated [CO2] relative to ambient [CO2] at timepoint 3. Fold changes in gene expression are displayed in a log2 scale.

**Fig. 12.**
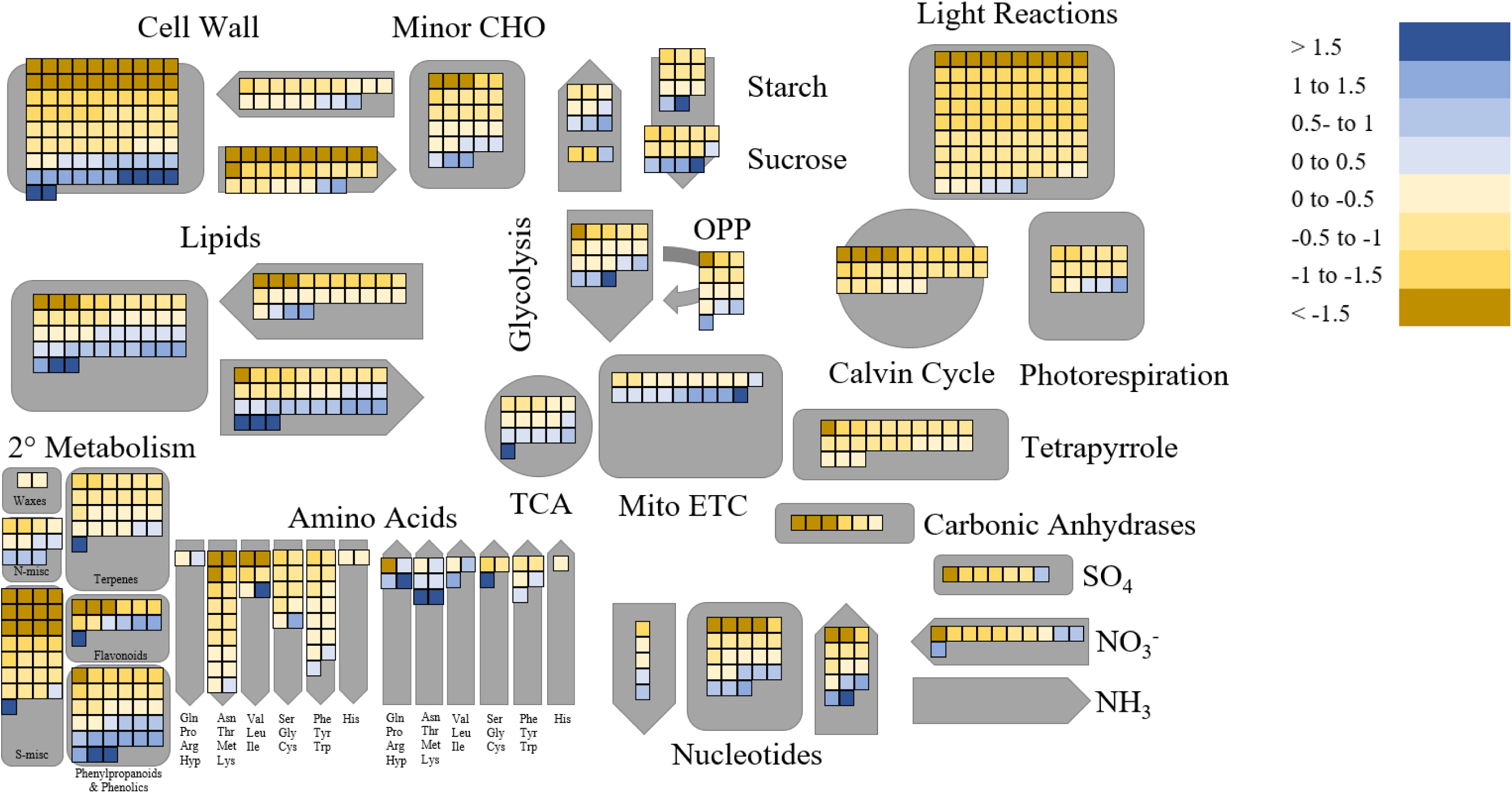
Transcript abundance in central metabolism pathways of *gnc* response to elevated [CO2] relative to ambient [CO2] at timepoint 3. Fold changes in gene expression are displayed in a log2 scale.

Contrasting the responses of *gnc* versus WT, the down-regulation of N metabolism genes at elevated [CO_2_] was stronger in *gnc* e.g. additional downregulation of nitrate reductase (NIA2), nitrite reductase (NIR1), and amino acid synthesis pathways. In addition, upregulation of respiration at elevated [CO_2_], especially mitochondrial electron transport, was greater in *gnc* than WT. At elevated [CO_2_], cell wall degradation and S-miscellaneous were down-regulated more, terpene metabolism was down-regulated less, and lipid degradation was up-regulated less in *gnc* than WT.

In transfer [CO_2_] compared to ambient [CO_2_], transcript abundance of genes involved in carbon, nitrogen, sulfur, secondary, cell wall and lipid metabolism were generally greater, with nucleotide synthesis and S-miscellaneous secondary metabolic pathways being the notable exceptions where transcript abundance was lower (Fig. 13 and 14). Upregulation of genes involved in sulfur metabolism was stronger in *gnc* plants compared to WT. While many aspects of this transcriptional response were again shared by *gnc* and WT, photosynthetic light reaction genes were upregulated in WT, but not in *gnc* plants.

**Fig. 13.**
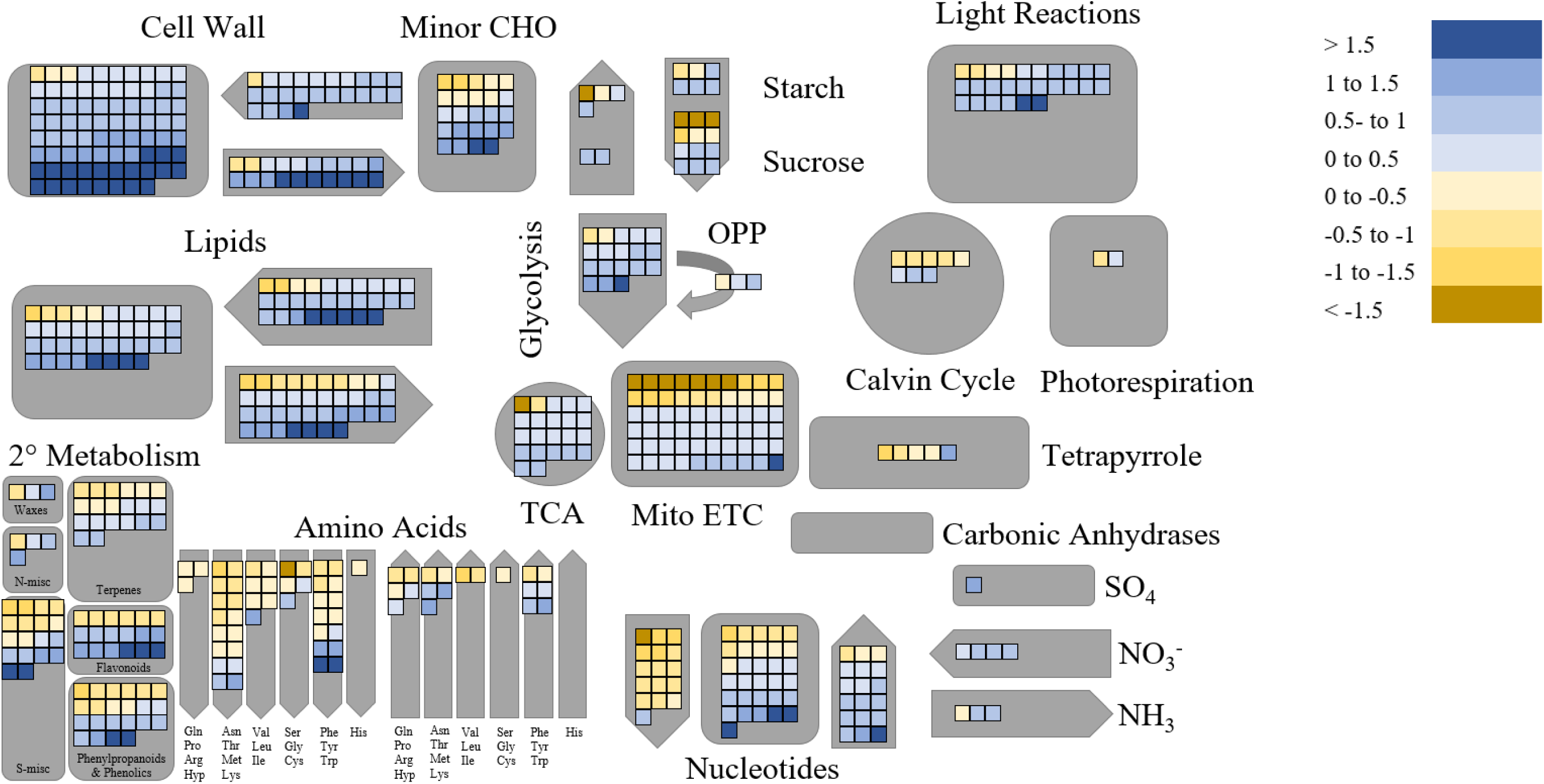
Transcript abundance in central metabolism pathways of WT response to transfer [CO2] relative to ambient [CO2] at timepoint 3. Fold changes in gene expression are displayed in a log2 scale.

### Knocking out GNC provokes the disruption of the network neighborhood around GNC

Coexpression networks were constructed to visualize changes in network topology. A correlation network (Pearson correlation coefficient |0.98|) of WT response to transfer [CO_2_] relative to ambient CO_2_ containing all of the first and second neighbors (1 hop) of GNC had 373 nodes and 7,629 edges with good connectivity (average number of neighbors = 40.906 or approximately 11%) (Fig. 15, Supplemental Table 4). This network construction approach captures both direct and indirect gene interactions with the GNC transcription factor. The characteristic path length between any two nodes was 2.445 and the network was moderately clustered with a coefficient of 0.536. While the network was not dense (0.11 density) and was not concentrated around a single node (0.195 centralization), it had a moderate tendency to contain hub nodes (0.614 heterogeneity) and was fully connected into a single component. The downregulation of GNC in mutants caused a topological disruption of the network where previously co-expressed genes no longer shared strong correlation coefficients resulting in a sparse network with few edges (220) between nodes (94) (Fig. 16, Supplemental Table 4).

**Fig. 14.**
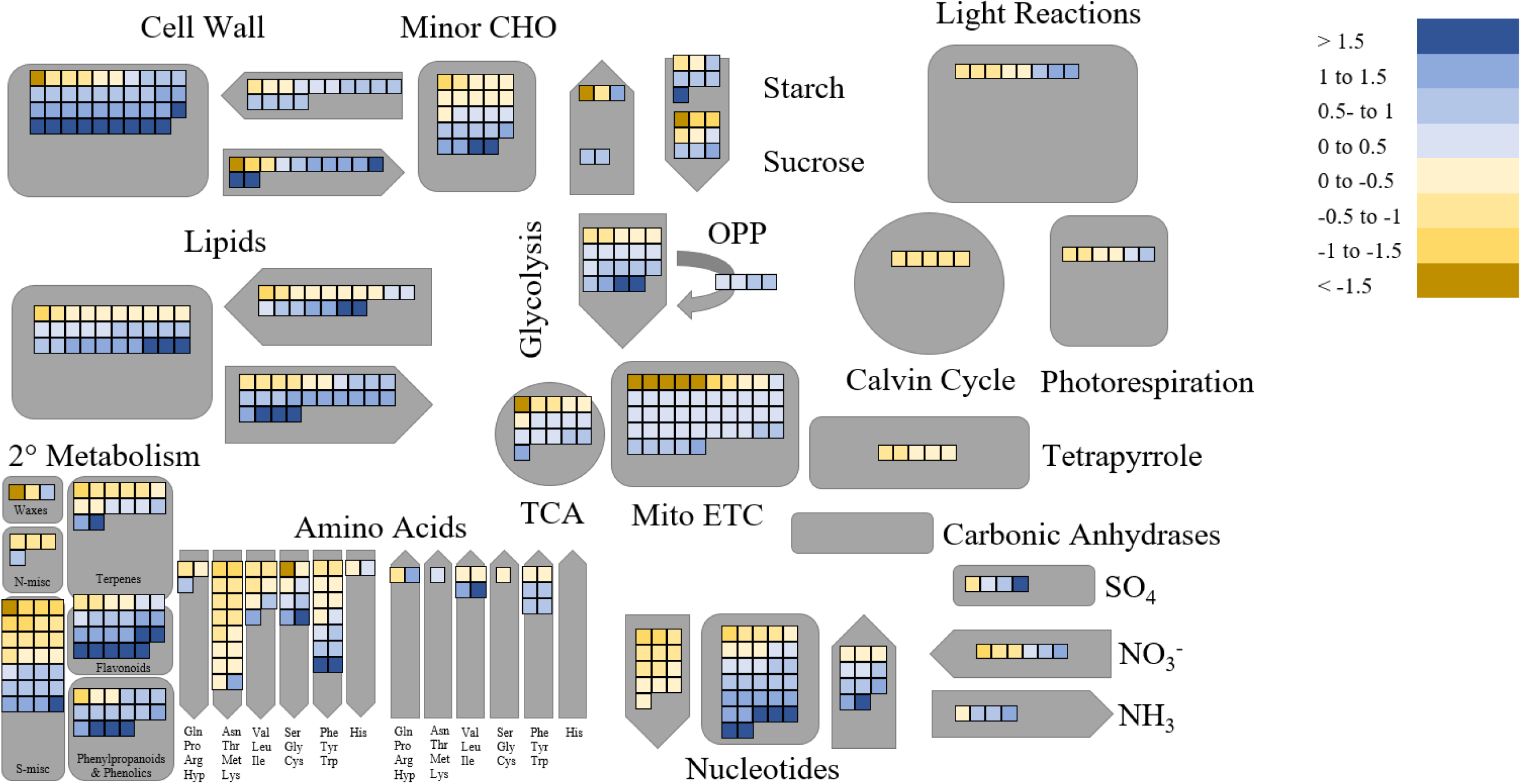
Transcript abundance in central metabolism pathways of *gnc* response to transfer [CO2] relative to ambient [CO2] at timepoint 3. Fold changes in gene expression are displayed in a log2 scale.

**Fig. 15.**
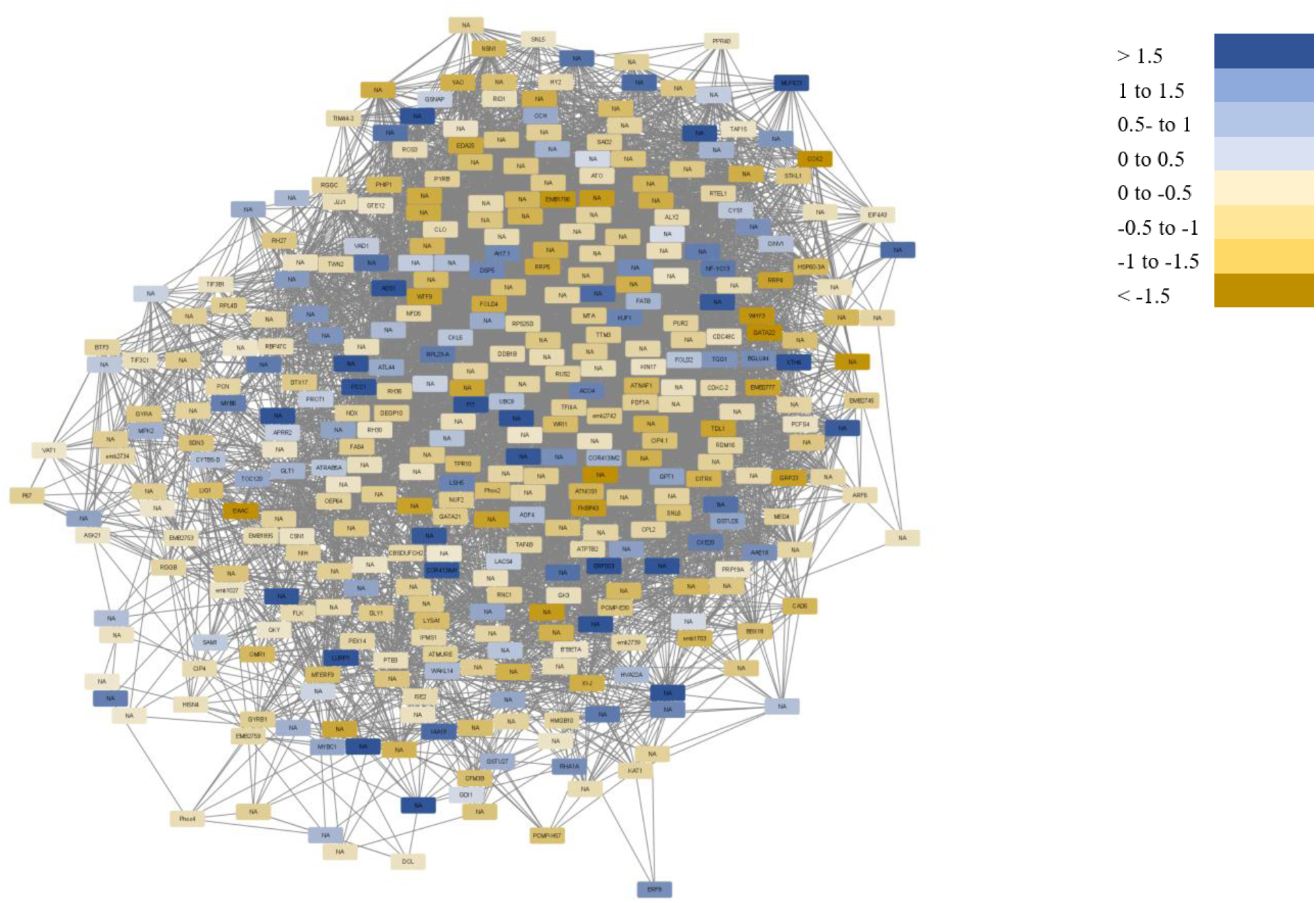
Unique correlation network of WT response to transfer [CO2] relative to ambient [CO2] including first and second neighbors (1 hop) of GNC.

**Fig. 16.**
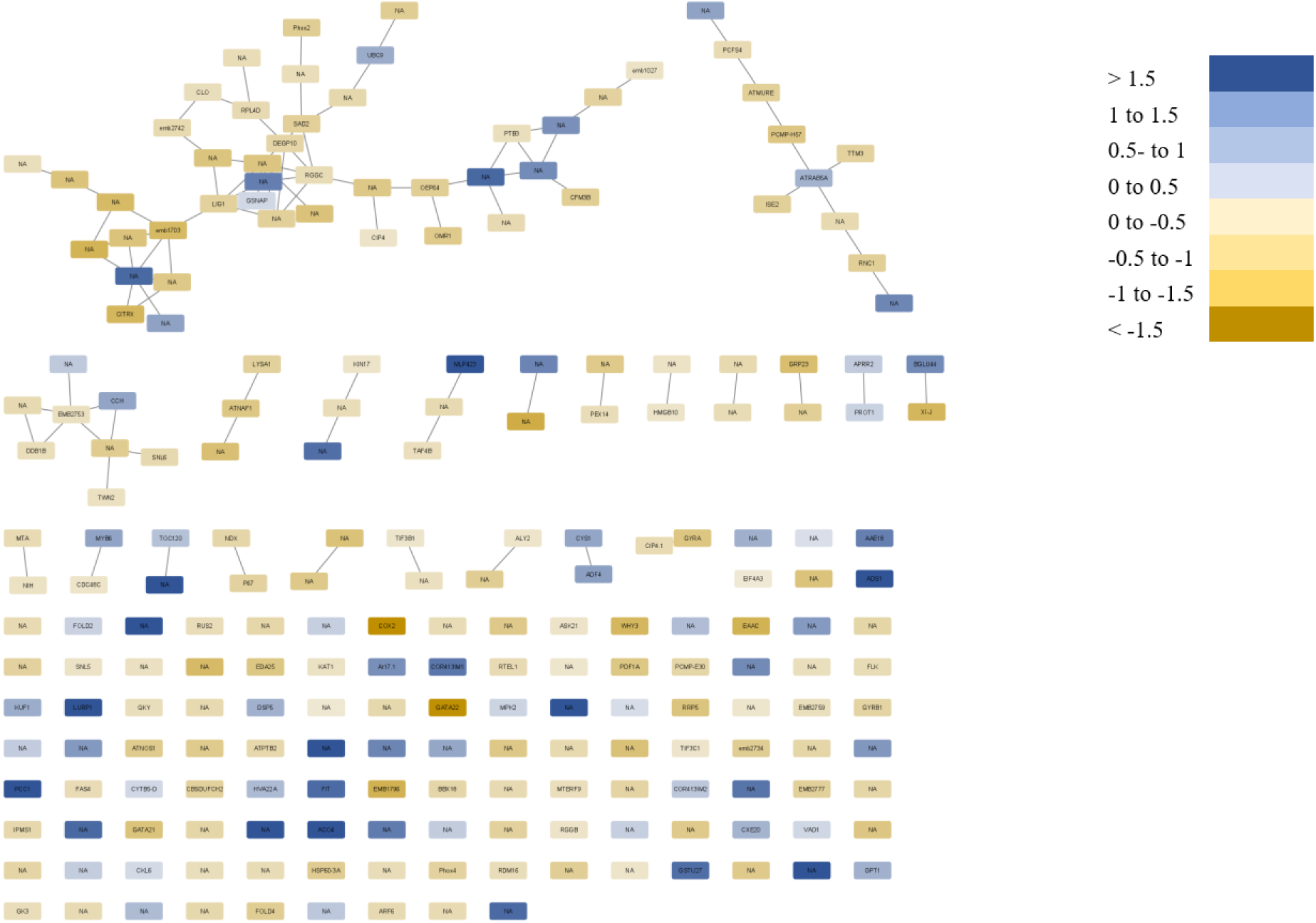
Unique correlation network of *gnc* response to transfer [CO2] relative to ambient [CO2] including first and second neighbors (1 hop) of GNC.

Pathways that were connected and no longer are include metabolic processes related to amino acids, amines, tRNA aminoacylation, carboxylic acids, oxoacids, organic acids, and ketones. Quantitatively, the network became less connected (average number of neighbors = 2.634 or approximately 1%) and less clustered (0.16 clustering coefficient). The distance between any two nodes also increased (characteristic path length= 4.954) and the network became fractured from a single module into 142 subnetworks.

This effect was exacerbated when networks were comprised of only first neighbors of GNC. The WT correlation network contained a decent number of nodes (38) and edges (291) with good connectivity (average number of neighbors = 15.316 or approximately 40%) (Supplemental Fig. 2, Supplemental Table 4). The characteristic path length between any two nodes was 1.586 and the network was fairly well clustered with a coefficient of 0.705. While the network was not very dense (0.414 density) and did not have a high tendency to contain hub nodes (0.445 heterogeneity), it had a moderate concentration of ties to a single node (0.619 centralization) and was fully connected into a single component. When GNC was knocked out, the correlation network was completely disrupted (Supplemental Fig. 3, Supplemental Table 4). Only 25 nodes remain, but have zero edges connecting them. Pathways that were connected and no longer are include metabolic processes related to tRNAs, non-coding RNAs, RNAs, amino acids, and amines.

## Discussion

This study identified GNC as a transcription factor that plays a role in regulating the strength of the CO_2_ fertilization effect on photosynthetic CO_2_ assimilation and plant biomass production. Specifically, knocking out expression of GNC prevented the significant stimulation of biomass production seen in WT exposed to both long-term elevated [CO_2_] and transfer from ambient [CO_2_] to elevated [CO_2_] late in vegetative development of Arabidopsis (Fig. 1). This is significant because very few regulatory genes have been implicated in plant metabolic and growth responses to elevated [CO_2_], even though adapting crops to exploit the greater availability of this key resource would have significant benefits (Ainsworth et al., 2008; Singer et al., 2020). The finding is novel despite the increasing availability of data describing plant transcriptional responses to elevated [CO_2_] as a source of candidate genes (Taylor et al., 2005; Ainsworth et al., 2006; Leakey et al., 2009a; Leakey et al., 2009b; Markelz et al., 2014a; Markelz et al., 2014b; Vicente et al., 2016; Vicente et al., 2019). Notably, the results of this study align with predictions from *in silico* analyses that members of the GATA transcription factor family are important in regulating CO_2_ response in soybean (Kannan et al., 2019). Additional novel insights resulted from exploring the transcriptomic and physiological responses of leaves after plants were transferred from ambient [CO_2_] to elevated [CO_2_] alongside the more typical analysis of plants grown long-term at ambient [CO_2_] and elevated [CO_2_]. This helped demonstrate that the response of *gnc* plants to the greater photoassimilate supply produced at elevated [CO_2_] is distinct from that of WT.

At ambient [CO_2_], knocking out expression of GNC in Arabidopsis had modest negative effects on V_cmax_ and J_max_ as measures of photosynthetic capacity (Fig. 3), such that photosynthetic light response curves of *gnc* and WT were indistinguishable at ambient [CO_2_] (Fig. 2). These observations, along with slightly lower chlorophyll content in *gnc*, and generally weak effects of knocking out GNC on carbon metabolism and biomass production under ambient [CO_2_], are consistent with previous studies of this gene (Bi et al., 2005; Richter et al., 2010; Hudson et al., 2011; Behringer et al., 2014) even though plants in this study received greater light supply and were studied at later stages of plant development. The overall N status of *gnc* plants grown at ambient [CO_2_] was also not significantly different from WT when assessed in terms of leaf N per unit area, leaf N per unit mass, or C:N ratio (Fig. 5). Minimal differences in the *gnc* plants compared to WT at ambient [CO_2_] are important because the effect of elevated [CO_2_] on carbon gain and growth is dependent on the size and vitality of plants. So, had strong pleiotropic effects of *gnc* on growth been observed under ambient [CO_2_], experiments focusing on variation in carbon metabolism responses to elevated [CO_2_] would have been impossible to interpret. Such challenges are widely recognized in research focused on improving plant tolerance to other abiotic factors and may explain why so much remains to be discovered about the regulatory factors controlling metabolic and growth responses to elevated [CO_2_].

Relative to ambient [CO_2_], physiological differences between *gnc* and WT were more apparent and more important in the treatments where plants grew at elevated [CO_2_] in the long-term (elevated [CO_2_] treatment) or were transferred from ambient [CO_2_] to elevated [CO_2_] 30 DAG (transfer [CO_2_] treatment). It has long been recognized that *A* is stimulated almost instantaneously when plants experience an increase in the [CO_2_] at which they are growing (Stitt, 1991). The magnitude of this initial stimulation of *A* is predictable based on V_cmax_ and J_max_, which describe the sensitivity of *A* to [CO_2_] at low and high [CO_2_], respectively (Long and Bernacchi, 2003). At the elevated [CO_2_] used in this study, photosynthesis is limited by J_max_. Therefore, the lower J_max_ of *gnc* relative to WT translated into lower *A* for *gnc* relative to WT in the transfer [CO_2_] and elevated [CO_2_] treatments. This means that some of the lower CO_2_ fertilization effect on biomass production in *gnc* versus WT was likely a product of the distinct physiology of the two genotypes at ambient [CO_2_]. Nonetheless, as described below, other observations in the experiment suggest *gnc* does respond to elevated [CO_2_] in a fundamentally distinct fashion, rather than simply displaying a muted version of the WT response. This contrasts with the situation observed when the strength of responses to elevated [CO_2_] are modulated by varying N supply (Markelz et al. 2014a). One example of the unique responses of *gnc* and WT is the comparison of the broad transcriptional responses to transfer [CO_2_] and elevated [CO_2_] treatments relative to ambient [CO_2_]. Three days into the transfer [CO_2_] treatment, 258 GO terms were over-represented with genes that were differentially expressed relative to ambient [CO_2_] (Supplemental Table 3). For the portions of the transcriptional response that were shared between *gnc* and WT, the magnitude of changes in transcript abundance were 15% weaker in *gnc* than WT (Fig. 10B). While this is consistent with the weaker initial stimulation of *A* by the transfer [CO_2_] treatment in *gnc* versus WT, 143 of the 256 GO terms were significantly responsive for only one genotype. Likewise, of the 207 GO terms that were over-represented with DE genes for elevated [CO_2_] versus ambient [CO_2_], 112 were significantly responsive for only one genotype (Supplemental Table 2).

In some growing conditions, and in some species, the stimulation of *A* by elevated [CO_2_] declines over the long-term (Stitt, 1991; Leakey et al., 2009a). This phenomenon is termed photosynthetic acclimation to elevated [CO_2_] and is interpreted to occur to optimize tissue stoichiometry for biomass production by re-allocating N from photosynthetic machinery to growth of new biomass when N supply or C sink size is limiting (Stitt, 1991; Long et al., 2004; Leakey et al., 2009a). Photosynthetic acclimation to elevated [CO_2_] was not observed in either *gnc* or WT, since there was no significant difference in *A*, V_cmax_, or J_max_ between transfer [CO_2_] and elevated [CO_2_] treatments (Figs 2, 3). This suggests that the role of GNC in regulating carbon and nitrogen metabolism in the current study must have been limited to pathways beyond photosynthesis that influence biomass production under elevated [CO_2_]. However, it is important to also note that the nitrogen supply used in this experiment was previously determined to be ample (Markelz et al., 2014a). If N supply was limiting, photosynthetic acclimation would be more likely to occur, and GNC could still have a role in regulating it. Also, significantly lower transcript abundance for a large number of photosynthetic genes was observed at elevated [CO_2_] versus ambient [CO_2_]. This is expected to be the molecular mechanism that is driven by sensing of carbohydrate accumulation at elevated [CO_2_] and triggers lower photosynthetic capacity (Moore et al., 1999). However, a number of other studies have also observed decreased expression of photosynthetic genes at elevated [CO_2_] under high nitrogen supply without a reduction in photosynthetic capacity occurring (Leakey et al., 2009b; Vicente et al., 2016). So, while decreased expression of photosynthetic genes may poise plants for photosynthetic acclimation at elevated [CO_2_], other steps must be needed to bring it to fruition. There are a wide array of post-transcriptional and post-translational mechanisms that could be explored as candidates to fill this gap in understanding (Mazzucotelli et al., 2008; Gray et al., 2020).

Nitrogen concentration is typically decreased by elevated [CO_2_] while C:N is generally greater due to dilution by greater carbon pools and changes in nitrogen acquisition and allocation (Stitt and Krapp, 1999; Ainsworth and Long, 2005). Elevated [CO_2_] can also significantly decrease shoot nitrogen assimilation (Bloom et al., 2010). In this experiment, we observed a decrease in leaf N per unit mass (i.e., N concentration) at both transfer [CO_2_] and elevated [CO_2_] that was statistically indistinguishable in *gnc* and WT. However, *gnc* and WT differed in SLA at ambient [CO_2_] and differed in how SLA responded to elevated [CO_2_] and transfer [CO_2_] treatments. This significant genotype by CO_2_ interaction was one of the strongest observed in the study. In WT, elevated [CO_2_] and transfer [CO_2_] decreased SLA relative to ambient [CO_2_], and did so equivalently (-22%, Fig. 7). This is consistent with many prior studies of plants at elevated [CO_2_] (Long et al., 2004). In contrast, SLA of *gnc* decreased less than WT in response to elevated [CO_2_] (-12%) and did not respond to transfer [CO_2_] at all. Ultimately, the combination of changes in SLA and leaf N per unit mass at elevated [CO_2_] led to significantly greater leaf N per unit area in WT but not *gnc*. Another measure of contrasting nitrogen status was significantly greater C:N at elevated [CO_2_] versus ambient [CO_2_] in *gnc* but not WT. This was not a consequence of greater dilution of N by additional carbohydrates at elevated [CO_2_] since *A* was stimulated less in *gnc* than WT. Rather, it implies that, when GNC is knocked out, mechanisms operating to adjust N acquisition or allocation are not operating normally, leading to greater imbalances in tissue stoichiometry than in WT. In a previous study of wheat grown at high N supply, the flag leaf content of most amino acids was reduced at elevated [CO_2_] compared to ambient [CO_2_] (Vicente et al. 2016). At elevated [CO_2_] in Arabidopsis, the decreased transcript abundance for genes synthesizing amino acids and greater transcript abundance degrading amino acids would drive such a change in metabolites. The greater strength of these responses in *gnc* than WT – in terms of additional genes (e.g., nitrate reductase and nitrite reductase) plus stronger changes in transcript abundance - may contribute to driving the greater leaf C:N that was observed. This notion is supported by a number of the amino acid synthesis and degradation genes being known binding targets of GNC (Xu et al., 2017), but further metabolomic analysis would be needed to provide confirmation.

In response to long-term, elevated [CO_2_], there were changes in transcript abundance for a significant number of biosynthetic pathways, including those involved in the metabolism of cell walls, lipids, and secondary metabolism. In the absence of metabolite data, strong conclusions cannot be drawn from this result. But again, *gnc* showed distinct responses from WT, particularly with respect to cell wall degradation, S-miscellaneous, terpene metabolism, and lipid degradation (Figs 11, 12). So, these may be contributing to changes in the turnover and content of metabolites that compose a large fraction of leaf biomass and could contribute to the observed changes in SLA, as well as being pathways to focus on in follow-up studies.

Focusing on the transfer [CO_2_] treatment, it is notable that little to no changes in transcript abundance were detected 1 hour and 25 hours after transfer of plants from ambient [CO_2_] to elevated [CO_2_], but thousands of transcripts had significantly altered abundance and coexpression 73 hours after the transfer (Table 1 and Fig. 15). This is consistent with the idea that the transcriptional responses to elevated [CO_2_] of the bulk leaf are driven by the changes in metabolism that elevated [CO_2_] biochemically triggers (Leakey et al., 2009b). If elevated [CO_2_] was directly sensed, as are changes in supply of N and light (Chen et al., 2004; Vidal et al., 2020), a rapid transcriptional response would be expected. However, if photoassimilate needs to accumulate at sustained concentrations beyond some threshold to trigger transcriptional reprogramming of metabolism, it is conceivable that one to three days of photosynthetic CO_2_ assimilation being stimulated after transfer to elevated [CO_2_] would be needed. Likewise, it is not surprising that pathways which use photoassimilate were upregulated in response to the transfer [CO_2_] treatment i.e. glycolysis, citric acid cycle, cell wall, and lipid metabolism. Similarly, when *gnc* plants were transferred from ambient [CO_2_] to elevated [CO_2_], they displayed coexpression network rewiring compared to WT, most notably due to differential response of transcripts involved in sulfur starvation response. During sulfur starvation, the sulfur reduction pathway is upregulated and incorporation of sulfur into downstream products (i.e. glucosinolates, glutathiones, cysteine, and methionine) is downregulated (Kopriva, 2006). GNC is known to bind sulfate transporter 4.2, which was upregulated in *gnc* mutants (+0.93 log2FC), and GSH1, the rate limiting step in glutathione biosynthesis, which was downregulated in *gnc* mutants (-0.48 log2FC) (Xu et al., 2017). APS reductase and serine acetyltransferase were also upregulated in *gnc* plants (APR1 +0.99; APR3 +2.1; SAT3 +0.62; SAT4 +2.8 log2FC), consistent with sulfur deficiency (Zhang et al., 2004; Kopriva, 2006). Sulfur deficiency is associated with decreased nitrogen uptake, accumulation of free amino acids, reduced photosynthesis, a decrease in chlorophyll content, and reduced yield, which were all observed in *gnc* plants in this study (Terry, 1976; Burke et al., 1986; Dietz, 1989; Karmoker et al., 1991; Kastori et al., 2008; Abadie and Tcherkez, 2019). It is not surprising that a reduction in photosynthesis occurs during sulfur deficiency since sulfur is assimilated in the chloroplast and also requires ATP and NADPH. Increases in [CO_2_] only exacerbate these competing processes (Abadie and Tcherkez, 2019; Lulofs, 2019).

Less intuitively, there was an upregulation of photosynthetic genes due to transfer [CO_2_] in WT. Almost every other abiotic factor that stimulates photosynthetic carbon gain i.e. greater supply of N, light, and water within the ranges where they are limiting, also triggers acclimation responses resulting in greater photosynthetic capacity (Thornley, 1998; Evans and Poorter, 2001; Flexas et al., 2006). And, those factors vary over time and space in a way that places immediate selective pressures on plants. Sustained changes in atmospheric [CO_2_] have occurred almost exclusively on geological timescales until anthropogenic CO_2_ emissions accelerated in recent decades. So, we speculate that mechanisms initially sensing greater photoassimilate availability trigger greater investment in photosynthetic machinery. GNC is a candidate to be involved in this process, since knocking it out disrupted co-expression and significantly dampened the up-regulation of photosynthetic genes in the transfer [CO_2_] treatment. But, significant additional work would be required to test this hypothesis.

In summary, there are very few examples in the literature of regulatory genes that modulate plant metabolic and productivity responses to elevated [CO_2_]. In this study, *gnc* plants had significantly smaller CO_2_ fertilization effects on photosynthesis and biomass production than WT in both long-term elevated [CO_2_] and when transferred from ambient to elevated [CO_2_] 30 DAG. Some of this effect can be attributed to differences in photosynthetic capacity that had little to no influence on carbon gain at ambient [CO_2_]. But, *gnc* did not simply show a muted version of the WT response. Instead, *gnc* showed greater changes in leaf N status than WT, as well as altered patterns of response in the key leaf allometric trait of SLA. Divergence between gnc and WT was also observed in transcriptional responses to elevated [CO_2_] and transfer [CO_2_] treatments. In particular, knockout of GNC led to greater downregulation of transcripts involved in N metabolism and upregulation of transcripts involved in sulfur assimilation. Since GNL was upregulated in GNC mutants, it is important to point out that the effects observed in this work could be because GNC was knocked out or GNL was over-expressed. While additional work is required to reveal the mechanistic basis for the responses reported here, this work provides a case study of a transcription factor that regulated plant responses to elevated [CO_2_]. More specifically, it partially validates the predictions from *in silico* analysis suggesting that GATA transcription factors are good candidates to perform a regulatory role at elevated [CO_2_] and could be targets for engineering improved crop productivity in the future (Ainsworth et al., 2008; Kannan et al., 2019; Singer et al., 2020).

## Supporting information

Supplemental Data Set 1

Supplemental Data Set 2

Supplemental Data Set 3

Supplemental Data Set 4

## Supplemental tables

**Supplemental Table 1.** List of primers used for genotyping and qPCR.

**Supplemental Table 2.** Gene ontology enrichment results for ambient [CO_2_] vs. elevated [CO_2_].

**Supplemental Table 3.** Gene ontology enrichment results for ambient [CO_2_] vs. transfer [CO_2_].

**Supplemental Table 4.** Network statistics for ambient [CO_2_] vs. transfer [CO_2_].

## Supplemental figures

**Supplemental Figure 1.** Relative expression of GNC in WT (Col-0) and *gnc* T-DNA mutants measured by RT-qPCR (n=4).

**Supplemental Figure 2.** Unique correlation network of WT response to transfer [CO_2_] relative to ambient [CO_2_] including first neighbors of GNC.

**Supplemental Figure 3.** Unique correlation network of *gnc* response to transfer [CO_2_] relative to ambient [CO_2_] including first neighbors of GNC.

## Supplemental data

**Supplemental Data Set 1.** Differentially expressed genes of WT response to elevated [CO_2_] relative to ambient [CO_2_].

**Supplemental Data Set 2.** Differentially expressed genes of *gnc* response to elevated [CO_2_] relative to ambient [CO_2_].

**Supplemental Data Set 3.** Differentially expressed genes of WT response to transfer [CO_2_] relative to ambient [CO_2_].

**Supplemental Data Set 4.** Differentially expressed genes of *gnc* response to transfer [CO_2_] relative to ambient [CO_2_].

## Acknowledgments

We would like to thank Ryan Boyd for preliminary results. We thank Pauline Lemonnier, Aishwarya Kammala, Mike Masters, Tim Wertin, Brad Dalsing, and Jesse McGrath for help with sample collection and growth chamber operation. We thank Lisa Ainsworth for use of lab space and equipment. We thank Matthew Brooks for helpful discussion on the manuscript.

## Author Contributions

Authors JCQ, AMC, and ADBL contributed to the design of the research. JCQ and KK performed the experiments and data collection. JCQ conducted data processing and analysis. KK and AMC advised on sequencing and analyses. JCQ, AMC, and ADBL contributed to data interpretation and discussion. JCQ, AMC, and ADBL wrote the original draft of the manuscript. All authors have revised and approved the final manuscript.

## Data Availability

Primary data for this manuscript are available at: X. Sequence data from this article can be found in the NCBI Gene Expression Omnibus (GEO) data libraries under accession number X.

## References

Abadie C and Tcherkez G (2019) Plant sulfur metabolism is stimulated by photorespiration. Communications Biology 2: 379

Ainsworth EA, Long SP (2021) 30 years of free-air carbon dioxide enrichment (FACE): What have we learned about future crop productivity and its potential for adaptation? Glob Chang Biol 27: 27–49

Ainsworth EA, Long SP (2005) What have we learned from 15 years of free-air CO_2_ enrichment (FACE)? A meta-analytic review of the responses of photosynthesis, canopy properties and plant production to rising CO_2_. New Phytol 165: 351–372

Ainsworth EA, Rogers A, Leakey ADB (2008) Targets for crop biotechnology in a future high-CO_2_ and high-O_3_ world. Plant Physiol 147: 13–19

Ainsworth EA, Rogers A, Vodkin LO, Walter A, Schurr U (2006) The effects of elevated CO_2_ concentration on soybean gene expression. An analysis of growing and mature leaves. Plant Physiol 142: 135–147

An Y, Zhou Y, Han X, Shen C, Wang S, Liu C, Yin W, Xia X (2020) The GATA transcription factor GNC plays an important role in photosynthesis and growth in poplar. J Exp Bot 71: 1969–1984

Andrews S (2010) FastQC: a quality control tool for high throughput sequence data. Available at: https://www.bioinformatics.babraham.ac.uk/projects/fastqc/

Aoyama S, Reyes TH, Guglielminetti L, Lu Y, Morita Y, Sato T, Yamaguchi J (2014) Ubiquitin ligase ATL31 functions in leaf senescence in response to the balance between atmospheric CO _2_ and nitrogen availability in Arabidopsis. Plant Cell Physiol 55: 293–305

Arp WJ (1991) Effects of source-sink relations on photosynthetic acclimation to elevated CO_2_. Plant, Cell Environ 14: 869–875

Bastakis E, Hedtke B, Klermund C, Grimm B, Schwechheimer C, Biology PS, München TU (2018) LLM-domain B-GATA transcription factors play multifaceted roles in controlling greening in Arabidopsis. Plant Cell 30: 582–599

Behringer C, Bastakis E, Ranftl QL, Mayer KFX, Schwechheimer C (2014) Functional diversification within the family of B-GATA transcription factors through the leucine-leucine-methionine domain. Plant Physiol 166: 293–305

Behringer C, Schwechheimer C (2015) B-GATA transcription factors – insights into their structure, regulation, and role in plant development. Front Plant Sci 6: 1–12

Bernacchi CJ, Singsaas EL, Pimentel C, Portis AR, Long SP (2001) Improved temperature response functions for models of Rubisco-limited photosynthesis. Plant Cell Environ 24: 253–259

Bi Y, Zhang Y, Signorelli T, Zhao R, Zhu T, Rothstein S (2005) Genetic analysis of Arabidopsis GATA transcription factor gene family reveals a nitrate-inducible member important for chlorophyll synthesis and glucose sensitivity. Plant J 44: 680–692

Bloom AJ, Burger M, Asensio JSR, Cousins AB (2010) Carbon dioxide enrichment inhibits nitrate assimilation in wheat and Arabidopsis. Science 328(5980): 899–903

Burke JJ, Holloway P, Dalling MJ (1986) The effect of sulfur deficiency on the organisation and photosynthetic capacity of wheat leaves. Journal of Plant Physiology 125(3-4): 371–375

Chen M, Chory J, Fankhauser C (2004) Light signal transduction in higher plants. Annu Rev Genet 38: 87–117

Cheng S, Moore B, Seemann JR (1998) Effects of short- and long-term Elevated CO_2_ on the expression of Ribulose-1, 5-Bisphosphate Carboxylase/ Oxygenase genes and carbohydrate accumulation in leaves of Arabidopsis thaliana (L.) Heynh. Plant Physiol 116: 715–723

Chiang Y, Zubo YO, Tapken W, Kim HJ, Lavanway AM, Howard L, Pilon M, Kieber JJ, Schaller GE, Sciences B, et al (2012) Functional characterization of the GATA transcription factors GNC and CGA1 reveals their key role in chloroplast development, growth, and division. Plant Physiol 160: 332–348

Czechowski T, Stitt M, Altmann T, Udvardi MK (2005) Genome-wide identification and testing of superior reference genes for transcript normalization. Genome Anal 139: 5–17

Dietz (1989) Leaf and chloroplast development in relation to nutrient availability. Journal of Plant Physiology 134(5): 544–550

Ekman A, Bulow L, Stymne S (2007) Elevated atmospheric CO_2_ concentration and diurnal cycle induce changes in lipid composition in Arabidopsis thaliana. New Phytol 174: 591– 599

Evans JR, Poorter H (2001) Photosynthetic acclimation of plants to growth irradiance: the relative importance of specific leaf area and nitrogen partitioning in maximizing carbon gain. Plant Cell Environ 24: 755–767

Ewels P, Magnusson M, Lundin S, Käller M (2016) MultiQC: summarize analysis results for multiple tools and samples in a single report. Bioinformatics 32(19): 3047–3048

Farquhar GD, Von Caemmerer S, Berry JA (1980) A biochemical model of photosynthetic CO_2_ assimilation in leaves of C3 species. Planta 90: 78–90

Flexas J, Bota J, Galme J, Medrano H, Ribas-Carbo M (2006) Keeping a positive carbon balance under adverse conditions : responses of photosynthesis and respiration to water stress. Physiol Plant 127: 343–352

Franklin KA, Ho S, Patel D, Kumar SV, Spartz AK, Gu C, Ye S (2011) Phytochome-interacting factor 4 (PIF4) regulates auxin biosynthesis at high temperature. Proc Natl Acad Sci 108: 20231–20235

Gasparini K, Costa LC, Brito FAL, Pimenta TM, Barcellos F, Araújo WL, Zsögön A, Ribeiro DM (2019) Elevated CO_2_ induces age-dependent restoration of growth and metabolism in gibberellin-deficient plants. Planta 250: 1147–1161

Ge Y, Guo B, Cai Y, Zhang H, Luo S (2018) Transcriptome analysis identifies differentially expressed genes in maize leaf tissues in response to elevated atmospheric [CO_2_]. J Plant Interact 13: 373–379

Gray SB, Kajala K, Rodriguez-medina J, Brady SM (2020) Translational regulation contributes to the elevated CO_2_ response in two Solanum species. 102(2): 383–397

Hanson J, Smeekens S (2009) Sugar perception and signaling — an update. Curr Opin Plant Biol 12: 562–567

Hewitt EJ, Smith TA (1975) *Plant Mineral Nutrition*. John Wiley and Sons, New York, NY, USA.

Huber W, Carey VJ, Gentleman R et al. (2015) Orchestrating high-throughput genomic analysis with Bioconductor. Nature Methods 12:115–121

Hudson D, Guevara D, Yaish MW, Hannam C, Long N, Joseph D, Bi Y, Rothstein SJ (2011) GNC and CGA1 modulate chlorophyll biosynthesis and Glutamate Synthase (GLU1 / Fd-GOGAT) expression in Arabidopsis. PLoS One. doi: 10.1371/journal.pone.0026765

Hudson D, Guevara DR, Hand AJ, Xu Z, Hao L, Chen X, Zhu T, Bi Y, Rothstein SJ (2013) Rice cytokinin GATA transcription factor1 regulates chloroplast development and plant architecture. Plant Physiol 162: 132–144

IPCC (2021) Summary for Policymakers. In: Climate Change 2021: The Physical Science Basis. Contribution of Working Group I to the Sixth Assessment Report of the Intergovernmental Panel on Climate Change [Masson-Delmotte, V., P. Zhai, A. Pirani, S.L. Connors, C. Péan, S. Berger, N. Caud, Y. Chen, L. Goldfarb, M.I. Gomis, M. Huang, K. Leitzell, E. Lonnoy, J.B.R. Matthews, T.K., Maycock, T. Waterfield, O. Yelekçi, R. Yu, and B. Zhou (eds.)].

Jang J, León P, Zhou L, Sheenl J (1997) Hexokinase as a sugar sensor in higher plants. Plant Cell 9: 5–19

Kang SG, Price J, Lin P, Hong JC, Jang J (2010) The Arabidopsis bZIP1 transcription factor is involved in sugar signaling, protein networking, and DNA binding. Mol Plant 3: 361–373

Kannan K, Wang Y, Lang M, Long SP, Marshall-colon A (2019) Combining gene network, metabolic, and leaf-level models show means to future-proof soybean photosynthesis under rising CO_2_. In silico Plants 1:1

Kaplan F, Kopka J, Sung DY, Zhao W, Popp M, Porat R, Guy CL(2007) Transcript and metabolite profiling during cold acclimation of Arabidopsis reveals an intricate relationship of cold-regulated gene expression with modifications in metabolite content. Plant J 50: 967– 981

Kaplan F, Zhao W, Richards JT, Wheeler RM, Guy CL, Levine LH (2012) Transcriptional and metabolic insights into the differential physiological responses of Arabidopsis to optimal and supraoptimal atmospheric CO_2_. PLoS One. doi: 10.1371/journal.pone.0043583

Karmoker JL, Clarkson DT, Saker LR, Rooney JM, Purves JV (1991) Sulphate deprivation depresses the transport of nitrogen to the xylem and the hydraulic conductivity of barley (Hordeum vulgare L.) roots. Planta 185: 269–278

Kastori R, Plesnicar M, Arsenijevic-Maksimovic I, Petrovic N, Pankovic D, Sakac Z (2008) Photosynthesis, chlorophyll fluorescence, and water relations in young sugar beet plants as affected by sulfur supply. Journal of Plant Nutrition 23(8): 1037–1049

Katari MS, Nowicki SD, Aceituno FF, Nero D, Kelfer J, Parnell Thompson L, Cabello JM, Davidson RS, Goldberg AP, Shasha DE, Coruzzi GM, Gutierrez RA (2010) Virtual Plant: A software platform to support systems biology research. Plant Physiol. 152: 500–515

Katny M, Hoffmann-Thoma G, Schrier A, Fangmeier A, Jager H, van Bel A (2005) Increase of photosynthesis and starch in potato under elevated CO_2_ is dependent on leaf age. J Plant Physiol 162: 429–438

Klermund C, Ranftl QL, Diener J, Bastakis E, Richter R, Schwechheimer C (2016) LLM-domain B-GATA transcription factors promote stomatal development downstream of light signaling pathways in Arabidopsis thaliana hypocotyls. Plant Cell 28: 646–660

Koch KE (1996) Carbohydrate-modulated gene expression in plants. Annu Rev Plant Physiol Plant Mol Biol 47: 509–540

Koini MA, Alvey L, Allen T, Tilley CA, Harberd NP, Whitelam GC, Franklin KA, Le L (2009) High temperature-mediated adaptations in plant architecture require the bHLH transcription factor PIF4. Curr Biol 19: 408–413

Kopriva S (2006) Regulation of sulfate assimilation in Arabidopsis and beyond. Annals of Botany 97: 479–495

Krapp A, Stitt M (1995) An evaluation of direct and indirect mechanisms for the “sink-regulation” of photosynthesis in spinach: changes in gas exchange, carbohydrates, metabolites, enzyme activities and steady-state transcript levels after cold-girdling source leaves. Planta 195: 313–323

Leakey ADB, Ainsworth EA, Bernacchi CJ, Rogers A, Long SP, Ort DR (2009a) Elevated CO_2_ effects on plant carbon, nitrogen, and water relations: six important lessons from FACE. J Exp Bot 60: 2859–2876

Leakey ADB, Xu F, Gillespie KM, McGrath JM, Ainsworth EA, Ort DR (2009b) Genomic basis for stimulated respiration by plants growing under elevated carbon dioxide. Proc Natl Acad Sci 106: 3597–3602

Li P, Ainsworth EA, Leakey AD, Ulanov A, Lozovaya V, Ort DR, Bohnert HJ (2008) Arabidopsis transcript and metabolite profiles: ecotype-specific responses to open-air elevated [CO_2_]. Plant Cell Environ 31: 1673–1687

Lichtenthaler HK, Wellburn AR (1983) Determinations of total carotenoids and chlorophylls *a* and *b* of leaf extracts in different solvents. Biochem Soc Trans 11(5): 591–592

Long SP, Ainsworth EA, Rogers A, Ort DR (2004) Rising atmospheric carbon dioxide: plants FACE the future. Annu Rev Plant Biol 55: 591–628

Long SP, Bernacchi CJ (2003) Gas exchange measurements, what can they tell us about the underlying limitations to photosynthesis? Procedures and sources of error. J Exp Bot 54: 2393–2401

Lulofs M (2019) The effects of elevated levels of CO_2_ on sulfur metabolism in Arabidopsis thaliana. [Bachelor’s thesis, University of Groningen]

Markelz RJC, Lai LX, Vosseler LN, Leakey ADB (2014a) Transcriptional reprogramming and stimulation of leaf respiration by elevated CO_2_ concentration is diminished, but not eliminated, under limiting nitrogen supply. Plant, Cell Environ 37: 886–898

Markelz RJC, Vosseller LN, Leakey ADB (2014b) Developmental stage specificity of transcriptional, biochemical and CO_2_ efflux responses of leaf dark respiration to growth of Arabidopsis thaliana at elevated [CO_2_]. Plant Cell Environ 37: 2542–2552

Mazzucotelli E, Mastrangelo AM, Crosatti C, Guerra D, Stanca AM, Cattivelli L (2008) Abiotic stress response in plants: When post-transcriptional and post-translational regulations control transcription. Plant Sci 174: 420–431

Miller R, Wu G, Deshpande RR, Vieler A, Gartner K, Li X, Moellering ER, Zauner S, Cornish AJ, Liu B, et al (2010) Changes in transcript abundance in chlamydomonas reinhardtii following nitrogen deprivation predict diversion of metabolism. Plant Physiol 154: 1737–1752

Moore B, Zhou L, Rolland F, Hall Q (2003) Role of the Arabidopsis glucose sensor HXK1 in nutrient, light, and hormonal signaling. 300: 332–337

Moore BD, Cheng S, Sims D, Seemann JR (1999) The biochemical and molecular basis for photosynthetic acclimation to elevated atmospheric CO_2_. Plant Cell Environ 22: 567–582

Nakashima K, Ito Y, Yamaguchi-shinozaki K (2009) Transcriptional regulatory networks in response to abiotic stresses in Arabidopsis and grasses 149: 88–95

Oelze M, Vogel MO, Alsharafa K, Kahmann U, Viehhauser A, Maurino VG, Dietz K (2012) Efficient acclimation of the chloroplast antioxidant defence of Arabidopsis thaliana leaves in response to a 10- or 100-fold light increment and the possible involvement of retrograde signals. J Exp Bot 63: 1297–1313

Ohnishi A, Wada H, Kobayashi K (2018) Improved photosynthesis in Arabidopsis roots by activation of GATA transcription factors. Photosynthetica 56: 1–12

Van Oosten J-J, Besford RT (1994) Sugar feeding mimics effect of acclimation to high CO_2_-rapid down regulation of RuBisCO small subunit transcripts but not of the large subunit transcripts. J Plant Physiol 143: 306–312

Para A, Li Y, Marshall-colón A, Varala K, Francoeur NJ, Moran TM (2014) Hit-and-run transcriptional control by bZIP1 mediates rapid nutrient signaling in Arabidopsis. Proc Natl Acad Sci 111: 10371–10376

Patro R, Duggal G, Love MI, Irizarry RA, Kingsford C (2017) Salmon provides fast and bias-aware quantification of transcript expression. Nature methods 14(4): 417

Paul MJ, Foyer CH (2001) Sink regulation of photosynthesis. J Exp Bot 52: 1383–1400

Penuelas J, Estiarte M (1998) Can elevated CO_2_ affect secondary metabolism and ecosystem function? TREE 13: 20–24

R Core Team (2017) R: A language and environment for statistical computing. R Foundation for Statistical Computing, Vienna, Austria. URL https://www.R-project.org/

Ranftl QL, Bastakis E, Klermund C, Schwechheimer C (2016) LLM domain containing B GATA factors control different aspects of cytokinin-regulated development in Arabidopsis thaliana. Plant Physiol 28: 646–660

Reyes C, Muro-pastor MI, Florencio FJ (2004) The GATA family of transcription factors in Arabidopsis and rice. Plant Physiol 134: 1718–1732

Richter R, Behringer C, Muller I, Schwechheimer C (2010) The GATA-type transcription factors GNC and GNL/ CGA1 repress gibberellin signaling downstream from DELLA proteins and PHYTOCHROME-INTERACTING FACTORS. Genes Dev 24: 2093–2104

Ritchie ME, Phipson B, Wu D, Hu Y, Law CW, Shi W, and Smyth GK (2015) limma powers differential expression analyses for RNA-sequencing and microarray studies. Nucleic Acids Research 43(7): e47

Robinson MD, Oshlack A (2010) A scaling normalization method for differential expression analysis of RNA-seq data. Genome Biology 11: R25

Robinson MD, McCarthy DJ, Smyth GK (2010) edgeR: a Bioconductor package for differential expression analysis of digital gene expression data. Bioinformatics 26(1): 139–140

Rolland F, Moore B, Sheen J (2002) Sugar sensing and signaling in plants. Plant Cell S185– S206

Sage RF, Sharkey TD, Seemann JR (1989) Acclimation of photosynthesis to elevated CO_2_ in five C3 species. Plant Physiol 89: 590–596

Saibo N, Lourenco T, Oliveira MM (2009) Transcription factors and regulation of photosynthetic and related metabolism under environmental stresses. Ann Bot 103: 609– 623

Sang Y, Sun W, Yang Z (2012) Signaling mechanisms integrating carbon and nitrogen utilization in plants. Front Biol 7: 548–556

Sato T, Maekawa S, Yasuda S, Sonoda Y, Katoh E, Ichikawa T, Nakazawa M (2009) CNI1 / ATL31, a RING-type ubiquitin ligase that functions in the carbon/ nitrogen response for growth phase transition in Arabidopsis seedlings. The Plant Journal 60(5): 852–864

Schmittgen TD, Livak KJ (2008) Analyzing real-time PCR data by the comparative CT method. Nat Protoc 3: 1101–1108

Shen C, Li Q, An Y, Zhou Y, Zhang Y, He F, Chen L, Liu C, Mao W, Wang X, Liang H, Yin W, Xia X (2022) The transcription factor GNC optimizes nitrogen use efficiency and growth by up-regulating the expression of nitrate uptake and assimilation genes in poplar. Journal of Experimental Botany 73(14): 4778–4792

Singer SD, Soolanayakanahally RY, Foroud NA, Kroebel R (2020) Biotechnological strategies for improved photosynthesis in a future of elevated atmospheric CO_2_. Planta 251: 1–28

Smith AM, Stitt M (2007) Coordination of carbon supply and plant growth. Plant, Cell Environ 30: 1126–1149

Stitt M (1991) Rising CO_2_ levels and their potential significance for carbon flow in photosynthetic cells. Plant Cell Environ 14: 741–762

Stitt M, Krapp A (1999) The interaction between elevated carbon dioxide and nitrogen nutrition: the physiological and molecular background. Plant, Cell Environ 22: 553–621

Soneson C, Love MI, Robinson MD (2015) Differential analyses for RNA-seq: transcript-level estimates improve gene-level inferences. F1000Research 4:1521

Talame V, Ozturk NZ, Bohnert HJ, Tuberosa R (2007) Barley transcript profiles under dehydration shock and drought stress treatments: a comparative analysis. J Exp Bot 58: 229–240

Taylor G, Street NR, Tricker PJ, Sjödin A, Graham L, Skogström O, Calfapietra C, Scarascia-mugnozza G, Jansson S (2005) The transcriptome of Populus in elevated CO_2_. New Phytol 167: 143–154

Teng N, Wang J, Chen T, Wu X, Wang Y, Lin J (2006) Elevated CO_2_ induces physiological, biochemical and structural changes in leaves of Arabidopsis thaliana. New Phytol 172(1): 92–103

Terry N (1976) Effects of sulfur on the photosynthesis of intact leaves and isolated chloroplasts of sugar beets. Plant Physiology 57: 477–479

Thomas PD, Ebert D, Muruganujan A, Mushayahama T, Albou L-P, Mi H (2021) PANTHER: Making genome-scale phylogenetics accessible to all. Protein Sci 31:8–22

Thornley JHM (1998) Dynamic model of leaf photosynthesis with acclimation to light and nitrogen. Ann Bot 81: 421–430

Torralbo F, Vicente R, Morcuende R, González-murua C, Aranjuelo I (2019) C and N metabolism in barley leaves and peduncles modulates responsiveness to changing CO_2_. J Exp Bot 70: 599–611

Usadel B, Bläsing OE, Gibon Y, Poree F, Höhne M, Günter M, Trethewey R, Kamlage B, Poorter H, Stitt M (2008) Multilevel genomic analysis of the response of transcripts, enzyme activities and metabolites in Arabidopsis rosettes to a progressive decrease of temperature in the non-freezing range. Plant Cell Environ 31: 518–547

Vicente R, Bolger AM, Martínez-carrasco R, Pérez P, Gutiérrez E, Usadel B, Morcuende R (2019) De novo transcriptome analysis of durum wheat flag leaves provides new insights into the regulatory response to elevated CO_2_ and high temperature. Front Plant Sci 10: 1–18

Vicente R, Perez P, Martinez-Carrasco R, Feil R, Lunn JE, Watanabe M, Arrivault S, Stitt M, Hoefgen R, Morcuende R (2016) Metabolic and transcriptional analysis of durum wheat responses to elevated CO_2_ at low and high nitrate supply. Plant Cell Physiol. 57(10): 2133–2146

Vidal EA, Alvarez JM, Araus V, Riveras E, Brooks MD, Krouk G (2020) Nitrate in 2020 : thirty years from transport to signaling networks. Plant Cell 32: 2094–2119

Wang P, Hendron R, Kelly S (2017) Transcriptional control of photosynthetic capacity : conservation and divergence from Arabidopsis to rice. New Phytol 216: 32–45

Xu Z, Casaretto JA, Bi Y-M, Rothstein SJ (2017) Genome-wide binding analysis of AtGNC and AtCGA1 demonstrates their cross-regulation and common and specific functions. Plant Direct 1–12

Zhang C, Hou Y, Hao Q, Chen H, Chen L (2015) Genome-wide Survey of the soybean GATA transcription factor gene family and expression analysis under low nitrogen stress. PLoS One 1–24

Zhang J, Du H, Chao M, Yin Z, Yang H, Li Y, Bender KW (2016) Identification of two bZIP transcription factors interacting with the promoter of soybean Rubisco Activase gene (GmRCAα). Front Plant Sci 7: 1–14

Zhang Z, Shrager J, Jain M, Chang C-W, Vallon O, Grossman AR (2004) Insights into the survival of chlamydomonas reinhardtii during sulfur starvation based on microarray analysis of gene expression. Eukaryotic Cell 3(5)

Zinta G, Abdelgawad H, Peshev D, Weedon JT (2018) Dynamics of metabolic responses to periods of combined heat and drought in Arabidopsis thaliana under ambient and elevated atmospheric CO_2_. J Exp Bot 69: 2159–2170

