## Supplementary figures and images for "GNC is a regulator of metabolic and productivity responses to elevated CO_2_ in *Arabidopsis thaliana*"

### Supplemental Fig 1

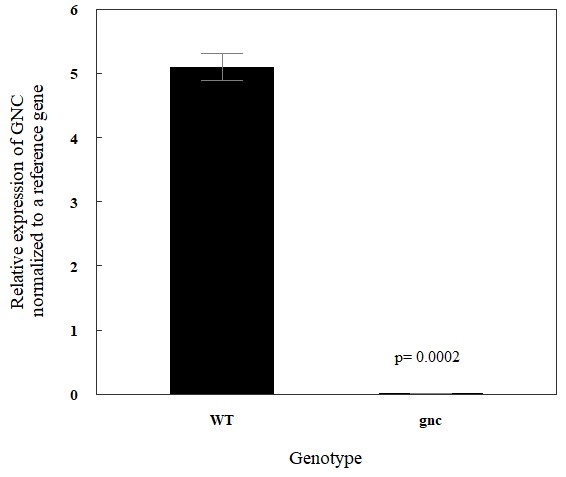

### Supplemental Fig 2

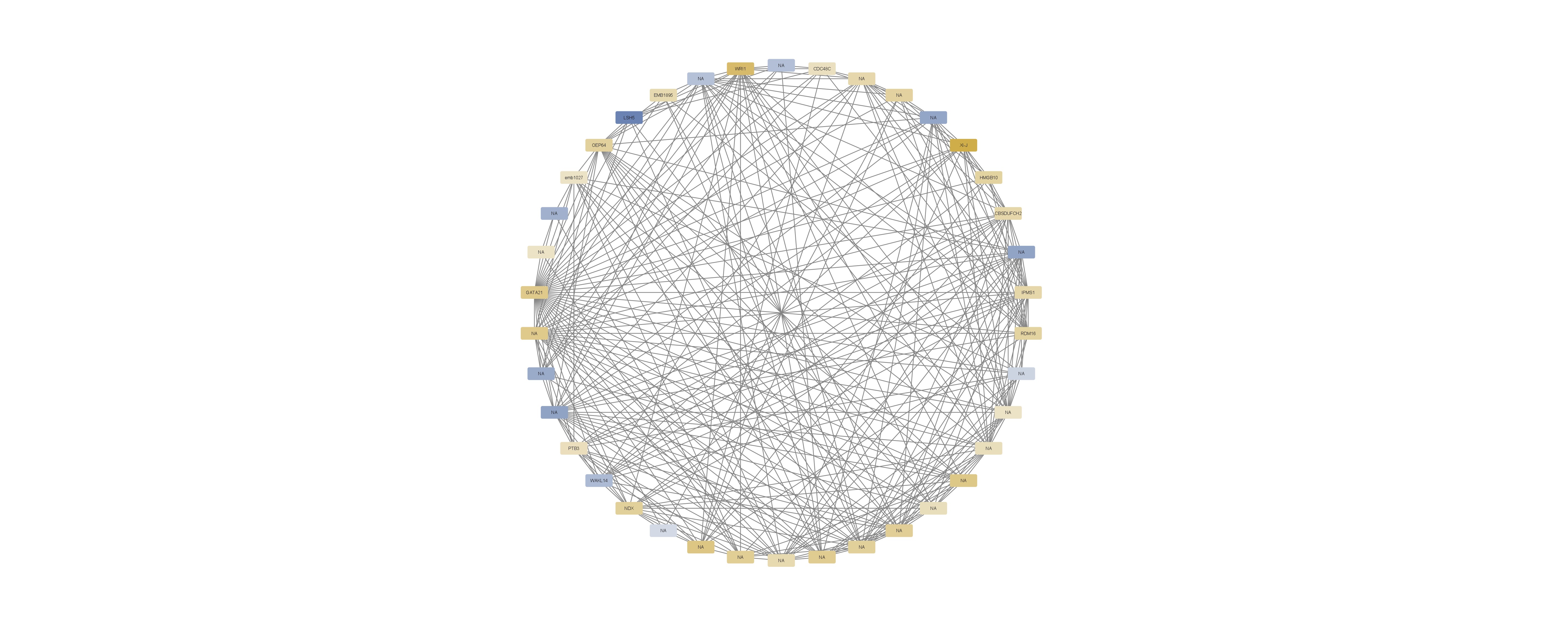

### Supplemental Fig 3

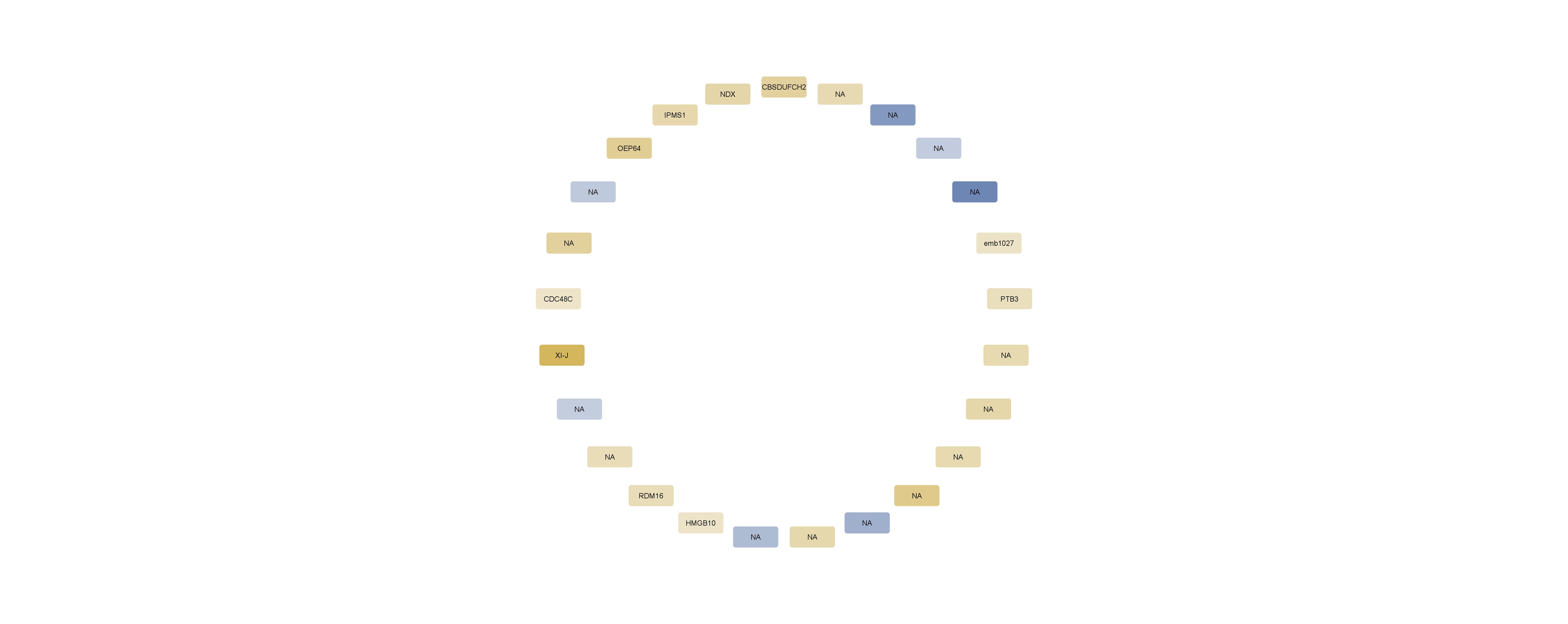
